# AtVPS13M1 regulates subcellular lipid distribution and lipid remodeling in phosphate-deprived plants

**DOI:** 10.1101/2024.05.22.594332

**Authors:** Sébastien Leterme, Catherine Albrieux, Sabine Brugière, Yohann Couté, Julien Dellinger, Benjamin Gillet, Sandrine Hughes, Julie Castets, Amélie Bernard, David Scheuring, Lucas Place, Marion Schilling, Juliette Jouhet, Morgane Michaud

**Affiliations:** Laboratoire de Physiologie Cellulaire et Végétale, CNRS, CEA, INRAE, Grenoble Alpes University, IRIG, CEA Grenoble, Grenoble, France; Univ. Grenoble Alpes, INSERM, CEA, UA13 BGE, CNRS, CEA, FR2048, 38000 Grenoble, France; Institut de Génomique Fonctionnelle de Lyon (IGFL), CNRS UMR 5242, Ecole Normale Supérieure de Lyon, Université Claude Bernard Lyon 1, Université de Lyon, Lyon, France; Laboratoire de Biogenèse Membranaire, UMR 5200, CNRS, Univ, Bordeaux, France; Department of Biology, University of Kaiserslautern-Landau, 67663 Kaiserslautern, Germany

## Abstract

VPS13 are conserved lipid transporters with multiple subcellular localizations playing key roles in many fundamental cellular processes. While the localizations and functions of VPS13 have been extensively investigated in yeast and animals, little is known about their counterparts in plants, particularly regarding their role in stress response. Here, we characterized AtVPS13M1, one of the four VPS13 paralogs in *Arabidopsis thaliana*. We show that AtVPS13M1 is expressed in young dividing and vascular tissues as a single major splice variant. AtVPS13M1 binds and transports glycerolipids with a low specificity *in vitro* and interferes with phospholipid metabolism in response to phosphate starvation, a nutrient stress that triggers a massive remodeling of membrane lipids. AtVPS13M1 is mainly located at the surface of mitochondria but is also present in microsomes. Finally, we demonstrated that AtVPS13M1 regulates phospholipid distribution between these two compartments, preventing their over-degradation in mitochondria while promoting their mobilization from microsomes. Overall, our findings highlight the conserved lipid transport function of VPS13 in plants, reveal its importance for mitochondrial membrane homeostasis and nutrient stress adaptation, and provide new insights into lipid remodeling mechanisms.

## Introduction

Vacuolar protein sorting 13 (VPS13) proteins are large, evolutionarily conserved eukaryotic proteins, typically comprising more than 3,000 amino acid residues ^1–3^. Although their overall structure and length are conserved, the number of VPS13 paralogs varies considerably across eukaryotes, ranging from one in *Saccharomyces cerevisiae* (ScVPS13) to six in charophytes. Four paralogs are found in both humans (HsVPS13A-D) and *Arabidopsis thaliana* (AtVPS13M1, AtVPS13M2, AtVPS13S and AtVPS13X) ^1–3^. In yeast and humans, VPS13 proteins localize to a wide range of organellar membranes and membrane contact sites (MCSs) ^4–6^. This diverse subcellular distribution is consistent with their involvement in numerous cellular processes, including meiosis and sporulation, maintenance of the actin cytoskeleton and cell morphology, mitochondrial function, regulation of phosphoinositide levels, and autophagosome and acrosome biogenesis ^4–6^. The importance of VPS13 proteins for cellular homeostasis is further highlighted by the association of mutations in human VPS13 genes with neurodegenerative disorders and other diseases ^7–10^.

Consistent with their localization at multiple MCSs, extensive physiological, biochemical and structural evidence has established VPS13 proteins as mediators of non-vesicular lipid transport between membranes ^5,6^. VPS13 proteins form elongated, rod-like structures approximately 20-30 nm in length that can bridge adjacent membranes at MCSs ^11–14^. Structural studies have revealed that these rods contain a continuous hydrophobic groove formed by repeating β-groove (RBG) domains. Their β-sheet-rich architecture creates a hydrophobic environment capable of accommodating lipids and facilitating their transport between membranes ^1,12,13,15^. In addition to this elongated lipid transport channel, VPS13 proteins contain various domains inserted between RBG repeats that contribute to their subcellular targeting and regulation ^1^. Most VPS13 proteins share four canonical domains: an N-terminal Chorein_N domain, a VPS13 adaptor-binding (VAB) domain, an ATG2_C domain and a C-terminal pleckstrin homology (PH)-like domain (Fig. 1A). They also frequently contain additional, non-canonical domains that vary among species and paralogs and that may contribute to the regulation of their localization and function ^1^. This diversity of localization determinants makes it challenging to predict the subcellular distribution of VPS13 proteins based solely on their domain architecture or computational predictions.

**Figure 1.**
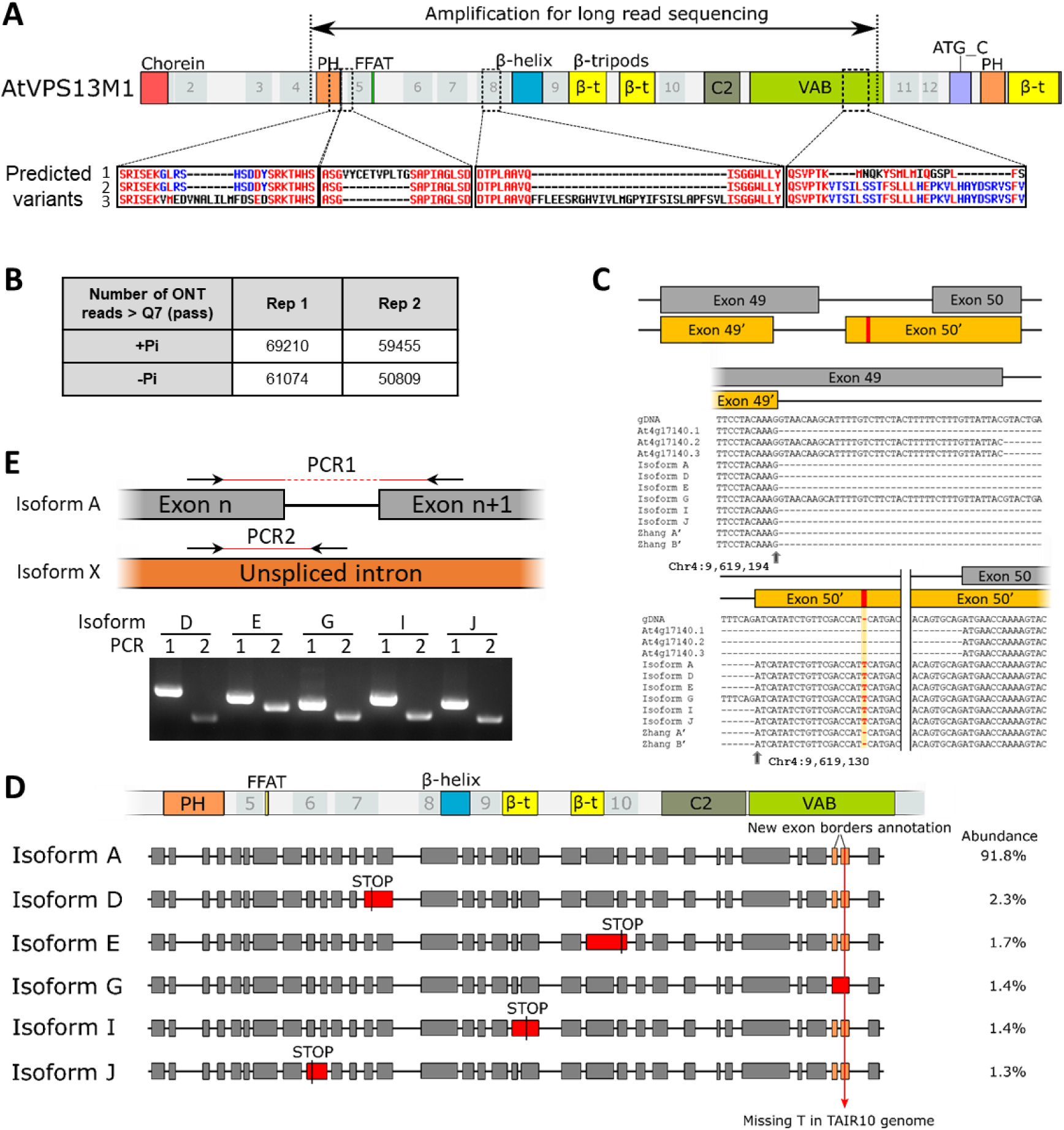
Long read sequencing analysis of AtVPS13M1 transcripts in calli grown in presence or absence of phosphate. **A.** Schematic representation of the AtVPS13M1 protein domains organization and position of the four predicted variable regions of At4g17140/AtVPS13M1 gene according to TAIR10 genome. The sequence alignments of the variable regions are shown. The range of the PCR performed for long read sequencing and spanning over 7600 bp is indicated above. The numbers represent the repeating β-groove (RBG) domains. Note that two of the predicted variable regions are located in domains presumably important for VPS13 protein localization (pleckstrin homology (PH)-like domain and VPS13 adaptor-binding (VAB) domain). **B.** Total number of Oxford Nanopore Technology (ONT) pass reads (>Q7) obtained for the two experimental replicates in + or in – phosphate (Pi) and used for the splicing variant analysis. **C.** Redefined borders of exons 49 and 50 of At4g17140/AtVPS13M1 gene. The original reference genome exon annotation is represented by grey boxes. The reannotated exons, here 49’ and 50’, are depicted in orange. The sequence alignments represent the right border of exon 49 and the left border of exon 50 according to the isoform A, D, E, G, I and J found by ONT long read sequencing, with precise positions on TAIR10 Chromosome 4 indicated (grey arrows). All sequences have been aligned on the reference gDNA. At4g17140.1, .2 and .3 are the three predicted splice variants cDNA sequences in TAIR10. Zhang A’ and Zhang B’ are the isoforms as described by Zhang *et al.* 2020. Notice the missing T base in the reference genome and in the two previously identified Zhang A’ and Zhang B’ sequences. **D.** Representation of the six AtVPS13M1 splice variants identified by long read sequencing and their relative abundance to each other. Grey squares indicate AtVPS13M1 exon distribution. The map of the corresponding protein domains is drawn on top. Red squares indicate the absence of intron splicing. Orange squares represent the newly annotated exons 49 and 50 with the correct exon borders. The red arrow represents the position of a missing T nucleotide in the TAIR10 genome. The abundance of the different identified isoforms is indicated on the right as a percentage over the six isoforms. **E.** Experimental validation by RT-PCR of the presence of the six AtVPS13M1 isoforms identified by ONT. Isoform D, E, G, J and I have a unspliced intron compared to isoform A and two PCR were performed to confirm their presence: PCR1, corresponding to amplification of the major isoform A, and PCR2, amplifying the unspliced intron in the corresponding

While VPS13 proteins have already been extensively studied in yeast and humans, their plant counterparts remain poorly characterized. VPS13 genes are present throughout Archaeplastida and have undergone multiple duplication, deletion and domain rearrangement events during the evolution of Viridiplantae ^3^. The *A. thaliana* genome encodes four VPS13 proteins, all of which contain the canonical VPS13 domains, except AtVPS13X, which lacks the C-terminal PH domain ^1^. Notably, AtVPS13M1 and AtVPS13M2 contain numerous additional non-canonical domains that, according to structural predictions, do not disrupt the overall folding of these proteins into elongated hydrophobic channel ^3^. AtVPS13M1 is the longest VPS13 protein in *A. thaliana*. It contains 12 RBG repeats, a FFAT motif potentially involved in endoplasmic reticulum (ER) targeting, and several additional domains, including an N-terminal PH domain, a C2 domain, a β-helix and three β-tripod domains (Fig. 1A) ^1,3^. Although the functions of the β-helix and β-tripod domains remain unknown, PH and C2 domains are known to mediate protein-protein and protein-lipid interactions. This complex domain architecture suggests that plant VPS13 proteins may exhibit diverse localization patterns and regulatory mechanisms, potentially reflecting plant-specific cellular requirements. To date, only a few physiological functions have been assigned to plant VPS13 proteins. AtVPS13S contributes to root growth, radial patterning and plant reproduction ^16,17^, whereas AtVPS13M2 was shown to promote pollen germination and pollen tube elongation by regulating vesicular trafficking at the pollen tube tip ^18^. A VPS13 protein has also been implicated in apomixis, a form of asexual plant reproduction producing seeds without meiosis or fertilization, in Taraxacum (dandelion) ^19^. However, the functions of other plant VPS13 proteins remain largely unexplored, and their lipid-binding properties and potential roles in non-vesicular lipid transport have not been yet investigated.

As sessile organisms, plants must adapt to a wide range of biotic and abiotic stresses, many of which trigger extensive changes in lipid homeostasis ^20,21^. Phosphate (Pi) starvation is a common nutritional stress that induces extensive remodeling of cellular membranes ^22–25^. Approximately one-third of intracellular Pi is stored in membrane phospholipids, which therefore constitute a major cellular Pi reservoir ^26^. Under Pi starvation, plants partially degrade their phospholipids to recycle the Pi contained in their polar headgroups. In parallel, digalactosyldiacylglycerol (DGDG), a phosphorus-free glycerolipid, is exported from chloroplasts to extraplastidial membranes, including the tonoplast, mitochondrial membranes and plasma membrane, where it partially replaces phospholipids to maintain membrane integrity ^23,27,28^. Although the metabolic enzymes involved in Pi starvation-induced lipid remodeling have been extensively characterized (reviewed in ^29,30^), the mechanisms mediating lipid transport between organelles remain poorly understood ^25^. In particular, only a few proteins involved in coordinating phospholipid degradation and DGDG redistribution have been identified. DGDG transport to mitochondria is thought to occur through non-vesicular pathways ^28^. The mitochondrial inner membrane protein AtMIC60 has been shown to regulate intramitochondrial DGDG import and phosphatidylethanolamine (PE) export ^31^. In addition, the chloroplast outer envelope protein LPTD1 participates in the import of precursors required for DGDG synthesis under Pi starvation ^32^.

To identify new components involved in lipid transport during Pi starvation, we investigated AtVPS13M1, a previously uncharacterized *A. thaliana* VPS13 paralog identified in mitochondrial outer membrane proteomic and complexomic analyses ^33,34^. We show that AtVPS13M1 is predominantly expressed in dividing and vascular tissues as a single major splice variant and has *in vitro* a broad lipid-binding capacity. We further demonstrate that AtVPS13M1 mediates lipid transport *in vitro* and contributes to phospholipid degradation during Pi starvation *in vivo*, supporting its function as a lipid transporter. AtVPS13M1 localizes to the mitochondrial surface as well as in microsomes, and contributes to phospholipid distribution between cell compartments. Together, our findings provide evidence that a plant VPS13 protein participates in the regulation of cell lipid homeostasis and Pi starvation-induced membrane remodeling. They also establish a foundation for investigating the contribution of VPS13 proteins to mitochondrial lipid homeostasis and the adaptation of plant membranes to nutrient limitation.

## Results

### Long-read sequencing reveals a single predominant AtVPS13M1 splice variant

To test whether phosphate (Pi) starvation affects AtVPS13M1 expression or splicing, we first quantified transcript levels by RT-qPCR. No significant differences were detected between Pi-replete and Pi-starved conditions (Fig. S1A). AtVPS13M1 (At4g17140) is a 23-kb gene with 63 exons, encoding a ∼13-kb transcript. The TAIR10 reference predicts three splice variants, differing in four regions that encode parts of the N-terminal PH and VAB domains that are critical for VPS13 localization (Fig. 1A). Thus, alternative splicing could alter AtVPS13M1 subcellular targeting. While two additional isoforms were previously identified by direct RNA sequencing ^35^, they were supported by only 3-4 reads, prompting us to comprehensively characterize AtVPS13M1 splice variants and assess their Pi-dependent regulation.

We performed Oxford Nanopore long-read sequencing of cDNA from *Arabidopsis thaliana* Col-0 calli grown 4 days under Pi-replete or Pi-starved conditions. As full-length AtVPS13M1 cDNA could not be amplified, we sequenced a 7.6-kb fragment spanning the four variable TAIR10 regions (Fig. 1A). After gel purification and filtering for read length and primer presence, we retained 50,809–69,210 high-quality reads per condition (Figs. 1B, S1B). Initial isoform identification using FLAIR ^36^ revealed discrepancies with TAIR10, including a single-nucleotide insertion at chromosome 4 position 9,619,112 and misannotated boundaries for exons 49-50 (Fig. 1C). After correcting the reference annotation with TALON ^37^, FLAIR identified 121 putative isoforms, 11 of which accounted for >1% of reads (Fig. S1C). Sequence analysis excluded five artifacts, leaving six splice variants (A, D, E, G, I, J) (Fig. 1D). Isoform A dominated, comprising >75% of reads, and served as the reference for comparisons. Importantly, Pi starvation did not alter isoform abundance or distribution (Fig. S1C).

All five alternative isoforms contained retained introns relative to isoform A. Only isoform G was predicted to encode a full-length protein (via an in-frame insertion), while the remaining four introduced premature termination codons (Fig. 1D). Isoforms A and D matched the two previously reported variants ^35^. To validate these findings, we performed isoform-specific PCR, designing primers to span retained introns or target intron-specific sequences (Fig. 1E). Amplification products confirmed all six exon-intron arrangements, validating the long-read sequencing results.

In summary, isoform A is the predominant AtVPS13M1 transcript, and neither its abundance nor alternative splicing is significantly affected by Pi starvation under the tested conditions.

### AtVPS13M1 is predominantly expressed in meristematic and vascular tissues

To map AtVPS13M1 protein tissue-specific expression, we generated a translational GUS reporter by inserting the *GUS* coding sequence into the *AtVPS13M1* genomic locus on a transformable artificial chromosome (TAC) ^38^. GUS was inserted after amino acid 472, a site structurally equivalent to a yeast *VPS13* insertion that preserves localization and function ^39^ (Fig. 2A). The construct, driven by the native *AtVPS13M1* promoter, was stably introduced into the *atvps13m1-3* mutant background (see below).

**Figure 2:**
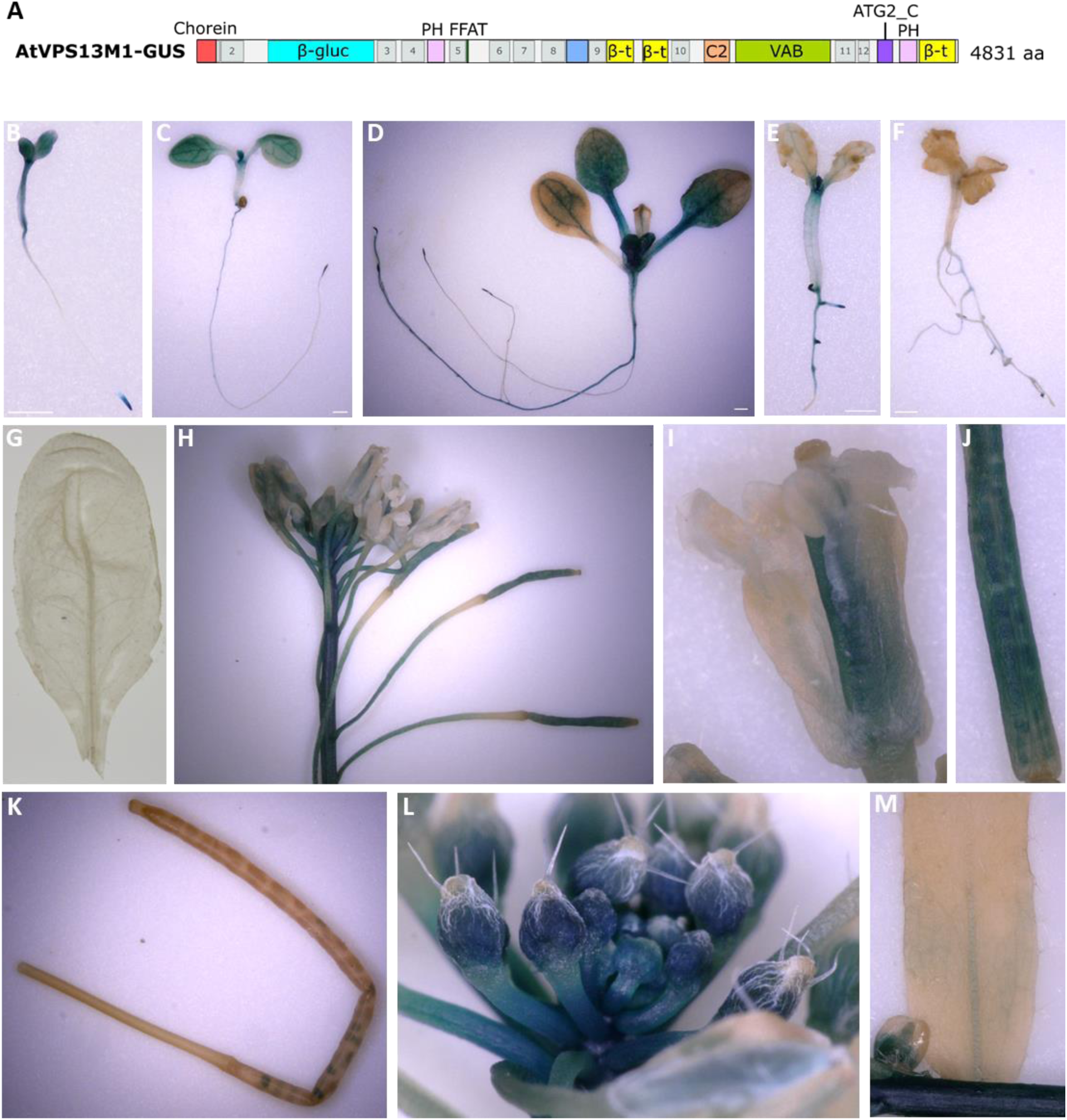
Analysis of AtVPS13M1 expression *in planta* in *Arabidopsis thaliana atvps13m1-3* KD stably expressing AtVPS13M1 fused to β-glucuronidase at position 472 within its native locus. Results from stably transformed line n°4 are shown. **A.** Schematic representation of the AtVPS13M1 protein fused to β-glucuronidase (β-gluc) used to analyze AtVPS13M1 expression *in planta*. **B-F.** Analysis of AtVPS13M1 expression by GUS staining in seedlings grown in liquid media containing (**B-D**) or not (**E-F**) phosphate (Pi). Seeds were germinated in a media containing Pi for three days (**B**) and then transferred in a media containing or not Pi and further grown for 3 (**C**: +Pi ; **E**: –Pi) and 6 days (**D**: +Pi ; **F**: –Pi). **G-M.** Analysis of AtVPS13M1 expression by GUS staining in leaf (**G**), inflorescence (**H**), flower (**I**), young (**J**) and mature (**K**) siliques, flower buds (**L**) and cauline leaf (**M**) from plants grown 7 weeks under long days conditions.

GUS staining of 3 to 9-day-old seedlings revealed strong AtVPS13M1 protein expression in root tips, lateral root initiation sites, and the shoot apical meristem, consistent with enrichment in actively dividing tissues (Figs. 2B-D, S2B-E). In roots, staining was absent in the elongation zone but detected in vascular tissues of mature regions (Figs. 2B, C, S2B). In cotyledons, staining was initially widespread at 3 days but became restricted to veins by 6 to 9 days post-germination (Figs. 2B-D, S2E). Consistent with this pattern, AtVPS13M1 protein was also detected in proliferating callus tissue (Fig. S2F).

To test whether Pi starvation alters AtVPS13M1 spatial distribution, 3-day-old seedlings were transferred to Pi-free medium for 3 or 6 days (Figs. 2E, F). As expected, Pi starvation inhibited primary root growth and promoted lateral root development ^40^. After 3 days of starvation, GUS staining disappeared from the primary root tip but persisted in developing lateral root tips, while cotyledon staining remained vein-restricted (Fig. 2C, E). After 6 days, seedling growth was severely inhibited, and GUS staining became barely detectable (Fig. 2D, F). These results suggest AtVPS13M1 protein distribution is tightly linked to tissue growth and developmental activity.

We next examined AtVPS13M1 distribution in 7-week-old plants (Figs. 2G-M, S2G-M). GUS staining was detected in inflorescence stems, floral buds, carpels, and young siliques (including valves and developing seeds) (Figs. 2G-J, S2G-L). In contrast, staining was reduced in mature siliques and restricted to a few basal, immature seeds (Figs. 2K, S2M). Weak staining was also observed at the base of cauline leaves, particularly in vascular tissues (Fig. 2M). No GUS staining was detected in rosette leaves, though confocal microscopy later revealed AtVPS13M1 fluorescence (see below), likely due to limited substrate penetration.

Together, these data show that AtVPS13M1 is widely expressed during plant development, with preferential accumulation in meristematic and vascular tissues. Its distribution shifts during Pi starvation and declines as tissues mature.

### AtVPS13M1 binds and transports lipids in vitro

VPS13 proteins are established lipid transporters. The N-terminal domain of yeast ScVPS13 binds and transports lipids *in vitro* ^41^. To test whether AtVPS13M1 shares this function, we expressed and purified a truncated form (AtVPS13M1(1-335)), which includes the N-terminal region homologous to lipid-transporting domains in yeast and mammalian VPS13 ^41,42^, fused to a C-terminal 6×His tag (Fig. S3A).

First, we assessed lipid binding capacity using nitrobenzoxadiazole (NBD)-labeled phospholipids (phosphatidylcholine (PC), phosphatidylethanolamine (PE), phosphatidylserine (PS) and phosphatidic acid (PA)) (Fig. 3A). After incubation with AtVPS13M1(1-335), Clear Native-PAGE and in-gel NBD fluorescence revealed robust lipid binding at the protein’s expected size, whereas no signal was detected for the negative control (Tom20.3 soluble domain) ^31^ (Fig. 3B). Thus, AtVPS13M1(1-335) binds multiple phospholipids, mirroring its yeast and human homologs.

**Figure 3.**
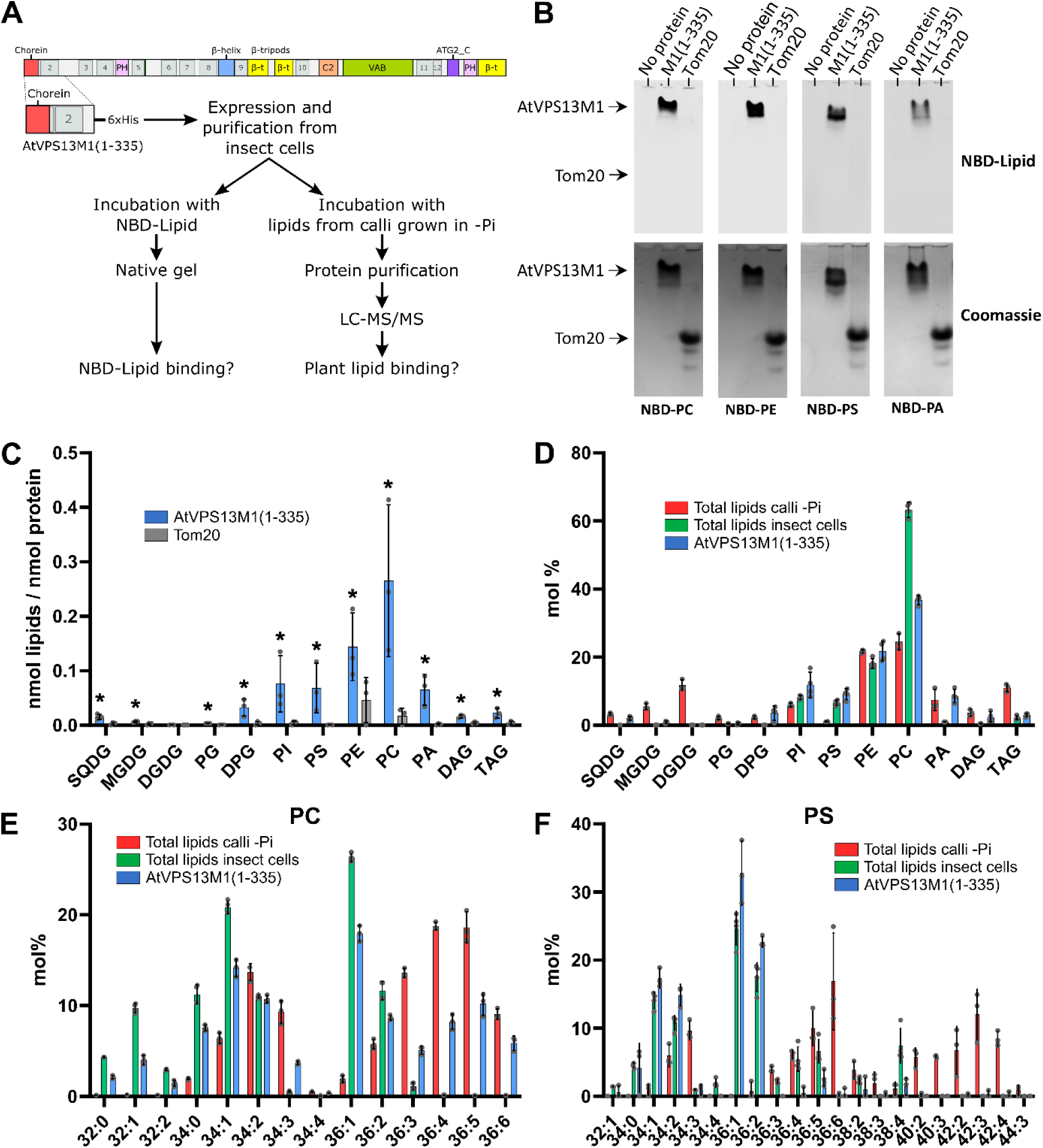
Analysis of AtVPS13M1 ability to bind lipids *in vitro*. **A.** Schematic representation of the experiments performed to analyze the ability of AtVPS13M1(1-335) fragment expressed and purified from insect cells to bind fluorescently labeled lipids (NBD-Lipid) or lipids from plant calli grown in absence of phosphate (Pi) *in vitro*. **B.** Binding analysis of AtVPS13M1(1-335) and Tom20 (negative control) to fluorescent lipids (NBD-PC, NBD-PE, NBD-PS or NBD-PA) on native PAGE. The upper and lower panels represent the NBD-Lipid fluorescence and protein Coomassie staining respectively. **C-D.** Mass spectrometry (MS) analysis of lipids bound to AtVPS13M1(1-335) or Tom20 (negative control) after incubation with calli total lipids and repurification. Results are expressed in nmol of lipids per nmol of proteins in order to analyze the absolute quantity of the different lipid classes bound to AtVPS13M1(1-335) compared to Tom20 negative control (**C**), and in mol% to assess the global distribution of lipid classes in AtVPS13M1(1-335) purifications compared to the total lipid extract of insect cells and calli (**D**). **E-F**. MS analysis of PC and PS lipid species in total lipids from calli, insect cells and AtVPS13M1 extracts. Results are expressed in mol%. **C-E.** Means, standard deviations and individual measures from each biological are represented, N=3. Statistical analysis was performed using non-parametric Mann-Whitney test, * p-value ≤ 0.05. DAG: diacylglycerol, DGDG: digalactosyldiacylglycerol, DPG: diphosphatidyldiacylglycerol, MGDG: monogalactosyldiacylglycerol, PA: phosphatidic acid, PC: phosphatidylcholine, PE: phosphatidylethanolamine, PG: phosphatidylglycerol, PI: phosphatidylinositol, PS: phosphatidylserine, SQDG: sulphoquinovosyldiacylglycerol, TAG: triacylglycerol.

Next, we tested binding capacity to physiological plant lipids, including plant-specific species (monogalactosyldiacylglycerol (MGDG), DGDG or sulfoquinovosyldiacylglycerol (SQDG)) never before evaluated for VPS13. AtVPS13M1(1-335) and Tom20.3 were incubated with lipid extracts from Pi-starved calli, followed by affinity purification and mass spectrometry quantification (Fig. 3A). AtVPS13M1(1-335) co-purified with a broad range of glycerolipids, whereas Tom20.3 showed minimal binding (Fig. 3C, D; Fig. S3B). PC and PE were the most abundant bound phospholipids, alongside neutral lipids (diacylglycerol (DAG) and triacylglycerol (TAG)) and mitochondria-specific diphosphatidyldiacylglycerol (DPG), hinting toward a link with mitochondria. Notably, AtVPS13M1(1-335) bound the chloroplastic lipids SQDG and MGDG but not DGDG, despite DGDG comprising >15% of the extract (Fig. 3D). Non-charged lipids (TAG, DAG) were underrepresented relative to total extracts, suggesting weak affinity (Fig. 3D).

To distinguish plant vs. insect cell-derived lipids, we analyzed their fatty acid saturation differences: insect lipids typically contain 0-1 unsaturations, while plant lipids can have up to 3. For MGDG, SQDG, and DPG, absent or undetectable in insect cells, only plant-derived species were bound, with no acyl-chain specificity (Fig. S3D, E, G). For other lipids (PC, PE, PI, PG, PA, DAG, TAG), AtVPS13M1(1-335) bound both plant and insect species (e.g., PC32:0 and PC32:2 from insect cells and PC36:4 and PC36:6 from plants) (Fig. 3E; Fig. S3C, F, H-K. PS was an exception: bound PS species (PS36:1, PS36:2) were exclusively insect-derived, while plant-specific PS (PS36:6, PS42:3) were absent. Overall, AtVPS13M1(1-335) binds diverse glycerolipids, excluding DGDG, without strong acyl-chain preference under these conditions.

To test lipid transport, we adapted a FRET-based assay (Kumar et al., 2018; Valverde et al., 2019) (Fig. 4A). Donor liposomes contained NBD-PS and Rhodamine (Rh)-PE and FRET quenching initially suppressed NBD fluorescence. AtVPS13M1(1-335), tethered via its His₆ tag, was added to donor liposomes. Without acceptor liposomes, a slight NBD fluorescence increase suggested lipid extrusion by AtVPS13M1(1-335) (Fig. 4C). Adding acceptor liposomes triggered a dose-dependent NBD fluorescence increase, indicating efficient lipid transfer from donor to acceptor liposomes (Fig. 4B, C). No signal increase occurred with Tom20.3 or without acceptor liposomes, confirming specificity (Fig. 4C). Finally, lipid transfer scaled with AtVPS13M1(1-335) concentration, directly linking transport activity to protein abundance (Fig. 4D).

**Figure 4.**
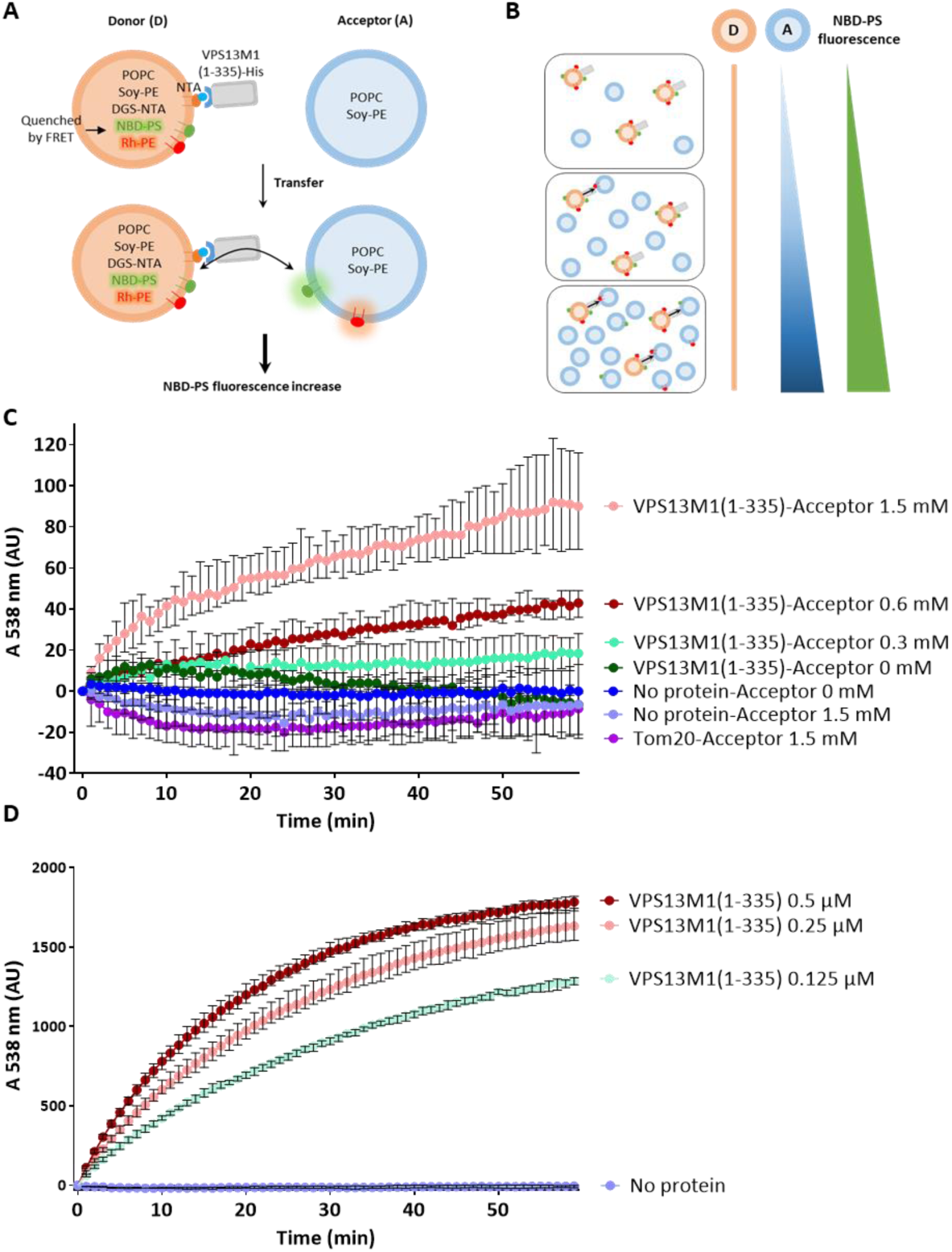
Analysis of AtVPS13M1 *in vitro* lipid transfer activity. **A-B**. Schematic representation of the *in vitro* fluorescent lipid transfer assay performed. Donor liposomes (25µM, 61% POPC, 30% soy PE, 2% NBD-PS, 2% rhodamine-PE and 5% DGS-NTA Ni) were incubated with or without AtVPS13M1(1-335) in the presence of acceptor liposomes (70% POPC, 30% soy PE). In this configuration, fluorescence of NBD-PS is quenched by rhodamine-PE by FRET. The transfer of fluorescent lipids triggers a dilution and a reduction of FRET signal, leading to an increase of NDB-PS fluorescence signal (**A**). As transfer requires proximity of donor and acceptor liposomes, the assay was performed at different acceptor liposomes concentrations to increase the proximity between liposomes and to promote the transfer (**B**). **C.** Transfer assay was performed with AtVPS13M1(1-335) at 0,125 µM in absence (Acceptor-0µM) or in presence of acceptor liposomes (A 0.3-1.5 mM). The transfer was monitored by following fluorescent signal increase at 538 nm for 60 min. AtVPS13M1(1-335) is attached to donor liposomes by a 6xHis tag in C-term. As a negative control, the experiment was performed with 0.125µM of Tom20.3 and 1.5 mM of acceptor liposomes. **D.** Donor liposomes were incubated with acceptor liposomes at 1.5 mM without or with different AtVPS13M1(1-335) concentrations. **C-D.** Medians with ranges are represented, **C**: N=4, **D**: N=3.

In summary, AtVPS13M1(1-335) binds a wide range of plant lipids and mediates lipid transport between close membranes *in vitro*.

### Phenotypic analysis of atvps13m1 mutants reveals no major roles in development, low-Pi growth, or autophagy

To investigate the physiological role of AtVPS13M1, we obtained four SALK T-DNA insertion lines ^43^ and identified homozygous plants (Figs. 5A, S4A, B). Surprisingly, RT-PCR showed that T-DNA insertions did not substantially reduce AtVPS13M1 transcript levels (Fig. S4C, D). Primers spanning the insertion sites confirmed that transcripts containing T-DNA were produced without apparent stability changes (Fig. S4C-E). As anti-AtVPS13M1 antibodies were unavailable, we assessed protein abundance by targeted mass spectrometry. While AtVPS13M1-specific peptides were detected in wild-type (Col-0) calli, none were detected in the four insertion lines (Fig S4F, G). Although peptide absence does not confirm complete protein loss, we designated these lines as AtVPS13M1-deficient mutants.

**Figure 5.**
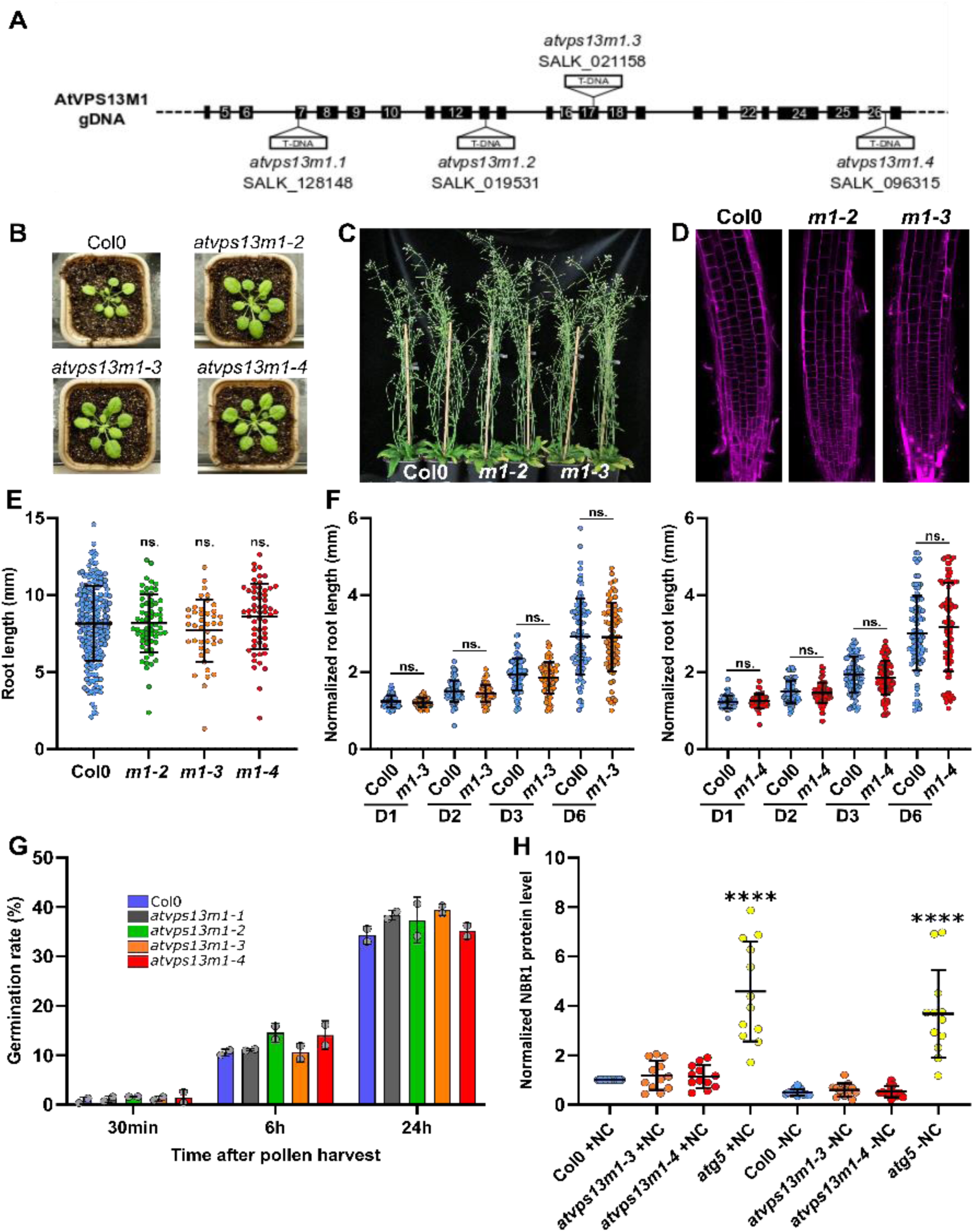
Plant phenotypic analysis of *atvps13m1 mutants*. **A.** Localization of the T-DNA inserted in the gDNA of the four *atvps13m1* mutants analyzed in this study. **B-C.** Analysis of Col0 and *atvps13m1* plants phenotype grown under long day conditions for 4 weeks (**B**) or 8 weeks (**C**) at 20/18°C. **D.** Analysis of root tip morphology by propidium iodide staining in 7 days-old Col0 and *atvps13m1-2* and *m1-3* mutant seedlings grown vertically. **E.** Root growth assays of Col0 and *atvps13m1* mutants grown vertically on long day conditions for 7 days. Root length (in mm) quantification in presented in the graph with the mean and SD. Col0: n=218, *m1-2*: n=63, *m1-3*: n=42, *m1-4*: n=57. **F.** Root growth assays of Col0 and *atvps13m1-3 and m1-4* mutants grown vertically on long day conditions for 4 days in 1 mM phosphate plates and transferred at day 0 (D0) in plates containing 5 µM of phosphate. Primary root length was measured at days 0 (D0), 1 (D1), 2 (D2), 3 (D3) and 6 (D6) after transfer. Root length (in mm) was normalized to the length at D0. Results are presented in the graph with the mean and SD. Col0 (left graph): n=80, *m1-3*: n=74, Col0 (right graph): n=93, *m1-4*: n=90. **G.** Analysis of pollen germination rate of Col0 and *atvps13m1* mutants. The results are expressed as percentage of germinated pollen 30min, 6h and 12h after harvesting. Means, standard deviations and individual measures are represented, N=2. **H.** Autophagy assessment by NBR1 protein level analysis by western blot in Col0, *atvps13m1-3* and *m1-4* mutant 7 days-old seedlings after 6h incubation in complete (+NC) or carbon and nitrogen starved (-NC) liquid media. The mutant *atg5* is used as a control of autophagy defect. Means, standard deviations and individual measure from independent biological replicates are represented, N=12.

To assess whether AtVPS13M1 contributes to plant growth or morphology, we grew Col0 and *atvps13m1-2* to *atvps13m1-4* lines on soil under standard conditions. No differences in growth rate or morphology were observed at any developmental stage (Fig. 5B, C).

Given that AtVPS13S (SHRUBBY), a VPS13 paralog in *A. thaliana*, causes poor root growth and defects in root cell layer organization ^16^, and that AtVPS13M1 is preferentially expressed in root tips (Fig. 2B, C), we investigated root morphology. Imaging of 7-day-old seedlings and propidium iodide staining of root cell walls revealed no substantial differences between Col0 and *atvps13m1-2* or *atvps13m1-3* (Fig. S5B-D). Root length measurements of 7-day-old seedlings grown on solid medium with 1 mM Pi also showed no significant differences (Fig. 5E), indicating AtVPS13M1 does not majorly influence root growth under standard Pi conditions.

We next tested primary root growth under low Pi. Col0, *atvps13m1-3*, and *atvps13m1-4* seeds were germinated for 4 days on 1 mM Pi plates and then transferred to 5 µM Pi. Primary root lengths were measured at 0, 1, 2, 3, and 6 days after transfer (Fig. 5F; Fig. S5D). Apart from a slight, non-significant reduction in root length at day 3, no major growth delay was observed in low Pi. We also examined pollen tube growth, as pollen tubes are among the fastest elongating plant structures and require extensive membrane transport ^44^. Pollen from Col0 and *atvps13m1-1* to *atvps13m1-4* lines was incubated in germination medium, and germination rates were determined after 30 min, 6 h, and 24 h (Fig. 5G). While germination rates increased over time for all lines, no differences were observed between Col0 and mutants, consistent with recent findings ^18^.

Finally, we investigated whether AtVPS13M1 plays a role in autophagy. In yeast, VPS13 and ATG2 proteins transport lipids to autophagosomal membranes to promote their growth ^45^. To test this, we monitored NBR1 protein levels, a receptor degraded during autophagy, in Col0, *atvps13m1-3*, *atvps13m1-4*, and the *atg5* autophagy-deficient mutant as a control. NBR1 levels were measured by Western blot in 7-day-old seedlings incubated for 6 h in either complete medium (+NC) or carbon- and nitrogen-starved medium (-NC) (Fig. S5E). As expected, NBR1 levels were elevated in *atg5* and did not decrease under -NC conditions (Fig. 5H). In contrast, NBR1 levels in *atvps13m1-3* and *atvps13m1-4* seedlings were similar to Col0 in both +NC and -NC conditions, suggesting AtVPS13M1 does not significantly impact autophagy induced by carbon and nitrogen starvation.

In summary, phenotypic analyses of *atvps13m1* mutants reveal no major roles for AtVPS13M1 in plant development, root growth under standard or low-Pi conditions, pollen tube growth, or autophagy.

### AtVPS13M1 contributes to phospholipid remodeling during Pi starvation

Given AtVPS13M1’s *in vitro* lipid transport activity, we tested whether its depletion alters lipid homeostasis during Pi starvation. We performed lipidomic analyses on non-photosynthetic, rapidly growing Arabidopsis calli, a model previously used to study mitochondrial lipid remodeling under Pi limitation ^31^. Wild-type and two independent mutant lines (*atvps13m1-1* and *atvps13m1-3*) were grown for 4 days under Pi-replete or Pi-starved conditions. Under Pi-replete conditions, mutants showed largely unchanged total glycerolipid abundance and composition, except for modest increases in MGDG and DGDG (*atvps13m1-3*) and PC (*atvps13m1-1*) (Fig. S6A-B).

Pi starvation reduced total glycerolipid abundance in all genotypes, but mutants retained significantly higher lipid levels than wild-type (Fig. 6A). This was accompanied by increased levels of several phospholipids (PG, PI, PC, PE) in both mutants, and a modest DGDG increase in *atvps13m1-3* (Fig. S6B). To quantify lipid remodeling, we calculated the log2 fold change (Log2FC) in lipid abundance under Pi-starved vs. Pi-replete conditions for each class, as well as for total phospholipids and galactolipids (Fig. 6B). This revealed a smaller reduction in PE abundance in both mutants and in total phospholipids in *atvps13m1-3*, with similar trends for PG and PC.

**Figure 6.**
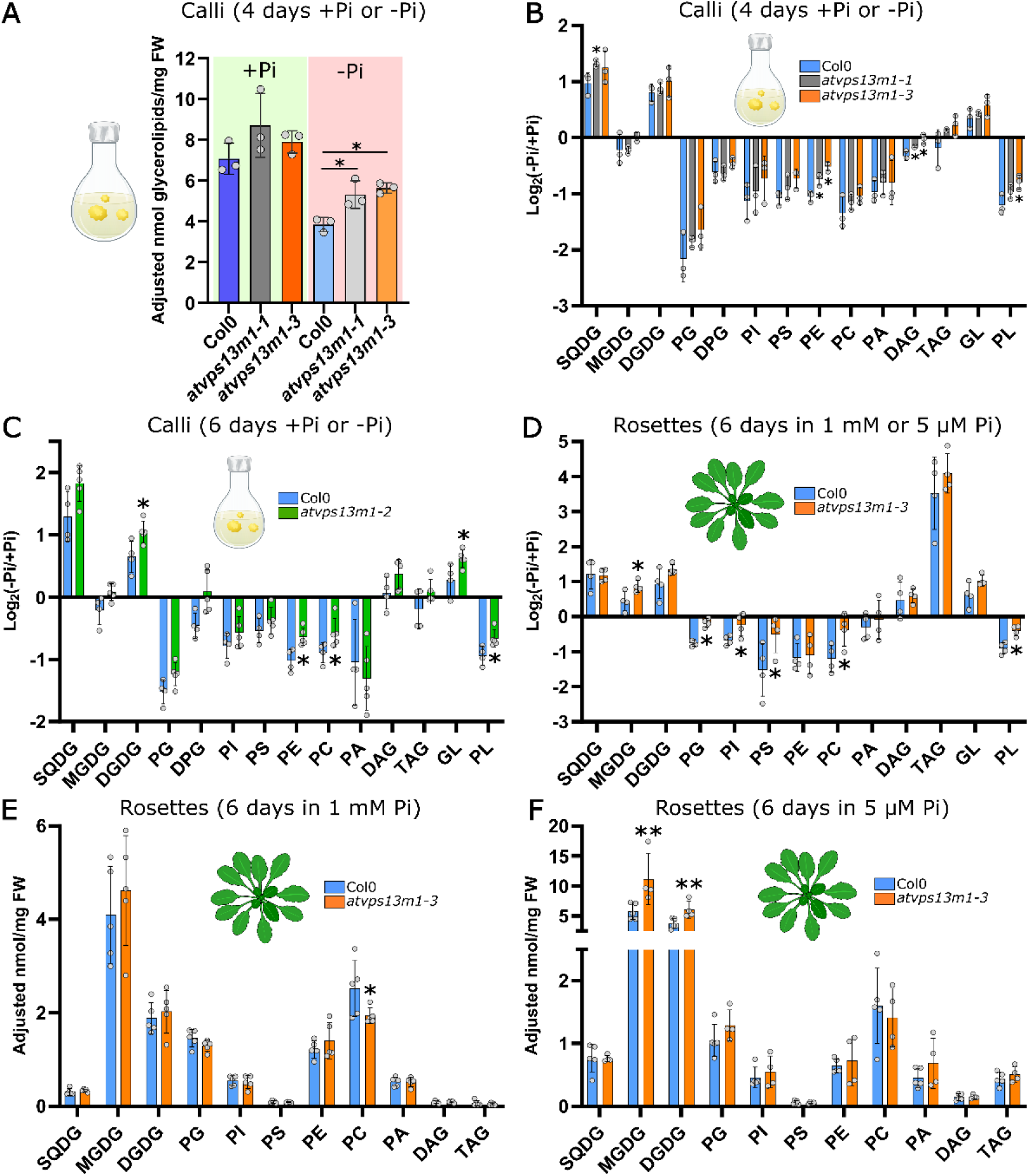
Lipid analysis in wild type (Col0) and *atvps13m1* calli or rosettes grown in presence and absence of phosphate (Pi). **A.** Adjusted total glycerolipid quantity (nmol) per mg of calli fresh weight in Col0 and *atvps13m1-1 and m1-3* mutants grown 4 days in presence or absence of Pi (see material and methods). **B.** Log_2_ fold change of each lipid class quantity obtained in -Pi versus +Pi in Col0 and *atvps13m1-1 and m1-3* mutants grown 4 days in presence or absence of Pi. **C.** Log_2_ fold change of each lipid class quantity obtained in -Pi versus +Pi in Col0 and *atvps13m1-2* mutants calli grown 6 days in presence or absence of Pi. **D.** Lipidomic analyses of Col0 and *atvps13m1-3* mutants rosettes grown 4 days in presence of 1 mM Pi and transferred 6 days in 1 mM or 5 µM Pi. Results are expressed as the Log_2_ fold change of each lipid species quantity obtained in 5 µM Pi versus 1 mM Pi conditions. **E-F**. Lipidomic analyses of Col0 and *atvps13m1-3* mutant rosettes grown 4 days in presence of 1 mM Pi and transferred 6 days in 1 mM or 5 µM Pi. Results are expressed in adjusted nmol of each lipid species per mg of rosettes fresh weight. Statistical analyses were performed using Mann-Whitney tests with a p-value ≤ 0,05 (*) or ≤0.01 (**). For all experiments, means, standard deviations and individual measures are represented, B-C: N=3, D: N=4, F-G: N=4-5. DAG: diacylglycerol, DGDG: digalactosyldiacylglycerol, DPG: diphosphatidyldiacylglycerol, GL: total galactoglycerolipids, MGDG: monogalactosyldiacylglycerol, PA: phosphatidic acid, PC: phosphatidylcholine, PE: phosphatidylethanolamine, PG: phosphatidylglycerol, PI: phosphatidylinositol, PL: total phospholipids, PS: phosphatidylserine, SQDG: sulphoquinovosyldiacylglycerol, TAG: triacylglycerol.

We validated these findings in a third mutant line, *atvps13m1-2*, after 6 days of Pi starvation (Figs. 6C, S6C-E). Consistently, mutant calli accumulated more glycerolipids than wild-type under Pi starvation (Fig. S6C), particularly DGDG, PE, and PC, though PE and PC differences were not statistically significant (Fig. S6D, E). Log2FC analysis confirmed a significantly smaller reduction in total phospholipids (especially PE and PC) and a greater increase in galactolipids in the mutant (Fig. 6C).

To test whether these alterations extend to photosynthetic tissues, we analyzed rosette lipid composition from wild-type and *atvps13m1-3* plants. Plants were grown in 1 mM Pi for 4 days, then transferred to Pi-replete (1 mM) or Pi-limiting (5 µM) medium for 6 days (Figs. 6D-F, S6F). Under low Pi conditions, mutant rosettes accumulated significantly more MGDG and DGDG than wild-type, resulting in higher total glycerolipid abundance (Figs. 6E-F, S6F). Although individual phospholipid classes did not differ significantly in abundance between genotypes under low Pi, Log2FC analysis revealed a smaller reduction in total phospholipids in the mutant, particularly of PG, PI, PS, and PC (Fig. 6D). Galactolipid abundance also tended to increase more strongly in the mutant.

Together, lipidomic analyses across three independent mutant lines and two tissue types show that AtVPS13M1 contributes to membrane lipid remodeling during Pi starvation. Its depletion consistently attenuates phospholipid reduction and enhances galactolipid accumulation, supporting a role in maintaining lipid homeostasis under Pi limitation.

### AtVPS13M1 localizes to mitochondrial surface and other organelles in A. thaliana

To determine the endogenous localization of AtVPS13M1, we introduced a 3xYepet fluorescent reporter at position 472 (AtVPS13M1-3xYepet, Fig. 6A) and stably expressed it in the *atvps13m1-3* background using the same recombination strategy as for GUS tagging (see above). Given the low expected abundance of VPS13 proteins ^46^, we used a triple Yepet tag to enhance signal detection. In stable lines, AtVPS13M1-3xYepet fluorescence was weak and often indistinguishable from background noise. However, the strongest signals were detected in vascular tissues of 6-week-old leaves, where Yepet fluorescence appeared as ∼1-µm-diameter round structures, potentially mitochondria (Fig. 7B, C; Fig. S7A, B). Similar structures were also observed in leaf epidermal cells, albeit at lower levels, indicating AtVPS13M1 expression is not restricted to vascular tissue (Fig. 7D; Fig. S7C).

**Figure 7:**
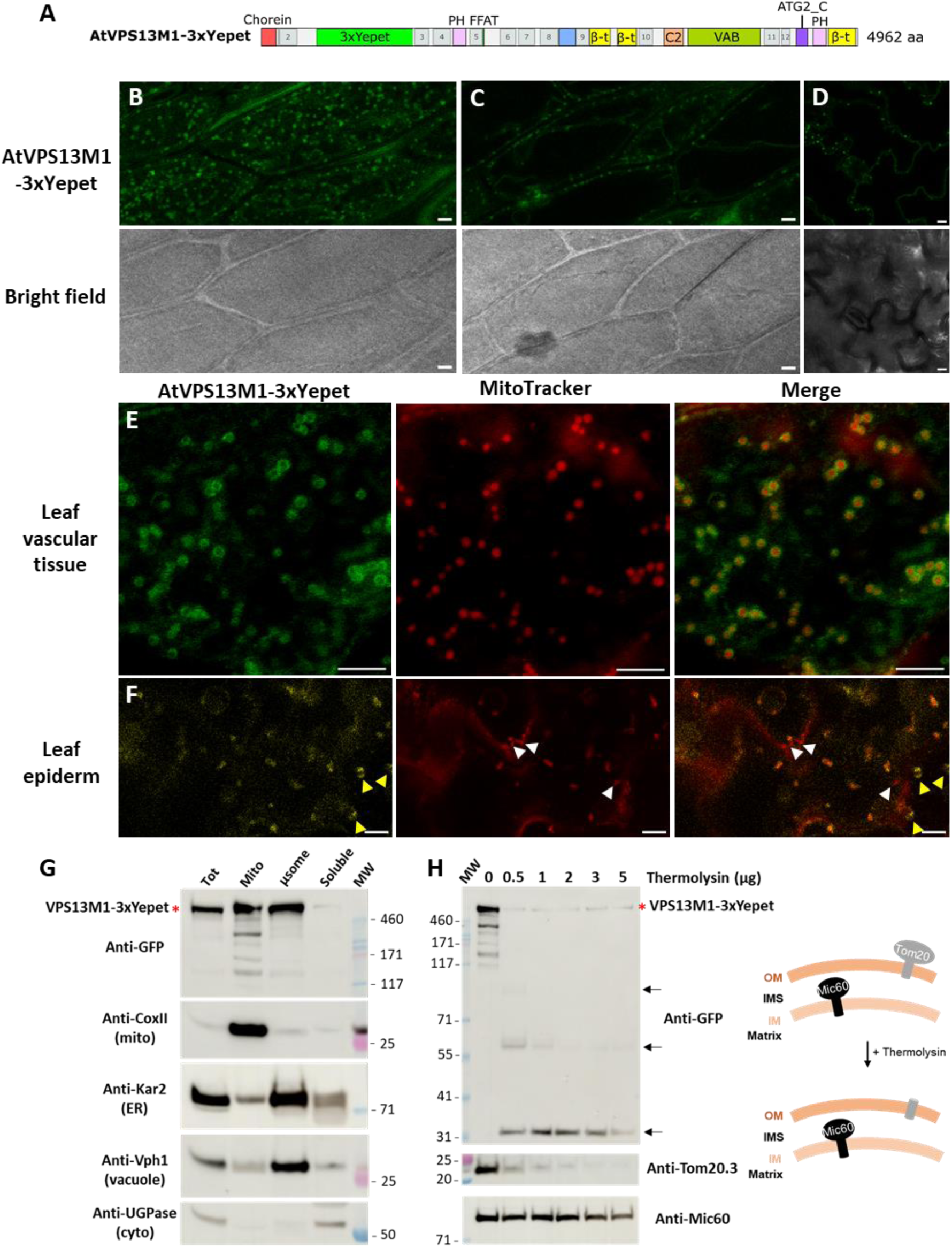
AtVPS13M1 subcellular localization in 6 weeks-old leaves or calli from *Arabidopsis thaliana atvps13m1.3* mutant stably expressing AtVPS13M1 fused to 3xYepet at position 472 expressed under its native promotor. **A-F**. Results from stably transformed line n°6 are shown. **A.** Schematic representation of the AtVPS13M1 protein fused to 3xYepet generated to analyze AtVPS13M1 subcellular localization *in planta.* The construction is under its genomic context (endogenous promoter and terminator, with introns). **B-C.** Localization in leaves vascular tissue at the cell cortex (**B**) or mid-plane (**C**). **D**. Localization in epidermal cells. **E-F.** Analysis of AtVPS13M1-3xYepet co-localization with mitochondria labeled with the Mitotracker^TM^ Red in leaf vascular (**E**) or epidermal (**F**) cells. Yellow arrows indicate AtVPS13M1-3xYepet signals which do not co-localized with Mitotracker^TM^ Red whereas white arrows indicate mitochondria which are not labeled by AtVPS13M1-3xYepet. **G.** Western blot analyses of subfractionation of Total (Tot), Mitochondria (Mito), Microsomes (µsome) and Soluble fractions from AtVPS13M1-3xYepet calli. **H.** Mitochondria from AtVPS13M1-3xYepet calli were incubated with increasing amount of thermolysin protease to analyze the sub-mitochondrial localization of AtVPS13M1. Arrow indicate AtVPS13M1 degradation products.

To test mitochondrial co-localization, we stained leaves with MitoTracker^TM^ Red FM. In vascular tissues, Yepet fluorescence formed ring-like structures around mitochondria, suggesting AtVPS13M1-3xYepet localizes to the mitochondrial surface (Fig. 7E). In epidermal cells, most Yepet signals co-localized with mitochondria, but some did not (Fig. 7F, white and yellow arrows), hinting at additional AtVPS13M1 localizations. Time-lapse imaging confirmed that AtVPS13M1-3xYepet remained associated with moving and fusing mitochondria, indicating stable mitochondrial localization (Fig. S7D). To verify specificity, we imaged non-transformed Col-0 leaves stained with MitoTracker^TM^ Red FM under identical confocal settings. Background fluorescence was rare and, when present, did not co-localize with mitochondria or resemble AtVPS13M1-3xYepet signals (Fig. S7E).

To confirm AtVPS13M1’s mitochondrial association biochemically, we performed cell fractionation on calli from a stable AtVPS13M1-3xYepet line. Western blots of total, mitochondrial, microsomal, and cytosolic fractions showed CoxII (mitochondrial), Kar2 (ER), Vph1 (vacuolar), and UGPase (cytosolic) markers were enriched in their respective fractions (Fig. 7G). In contrast, AtVPS13M1-3×Yepet was enriched in both mitochondrial and microsomal fractions, supporting its localization to mitochondria and other membrane structures. Its low cytosolic presence suggests AtVPS13M1 is primarily membrane-associated in calli.

To resolve its submitochondrial localization, we treated isolated mitochondria with increasing thermolysin concentrations (Fig. 7H). Like the outer membrane protein TOM20.3, AtVPS13M1 was highly sensitive to thermolysin, whereas the inner membrane protein AtMIC60 was protected. This confirms that AtVPS13M1 localizes to the mitochondrial surface.

### The absence of AtVPS13M1 perturbs phospholipid homeostasis in mitochondria and microsomes

Mitochondrial membranes undergo lipid remodeling during Pi starvation, as do vacuolar and plasma membranes ^27,28^. Given AtVPS13M1’s localization to both mitochondria and microsomes (Fig. 7G), we performed lipidomic analyses on whole seedlings, isolated mitochondria, and microsomes from Col0 and *atvps13m1-3* seedlings. Plants were grown for 4 days in 1 mM Pi and then transferred for 4 days to media containing 1 mM or 0 mM Pi. In whole seedlings and microsomes, the mutant had lower total lipid levels than Col0 under Pi-replete conditions (1 mM Pi) (Fig. 8A). Under Pi starvation (0 mM Pi), total lipid levels significantly decreased in Col0 but not in the mutant, suggesting impaired lipid degradation, consistent with our previous observations in calli and rosettes (Fig. 6, Fig. S6). In contrast, mitochondria in the mutant showed the opposite trend: lipid accumulation under Pi-replete conditions and a significant lipid decrease under Pi starvation (Fig. 8A).

**Figure 8.**
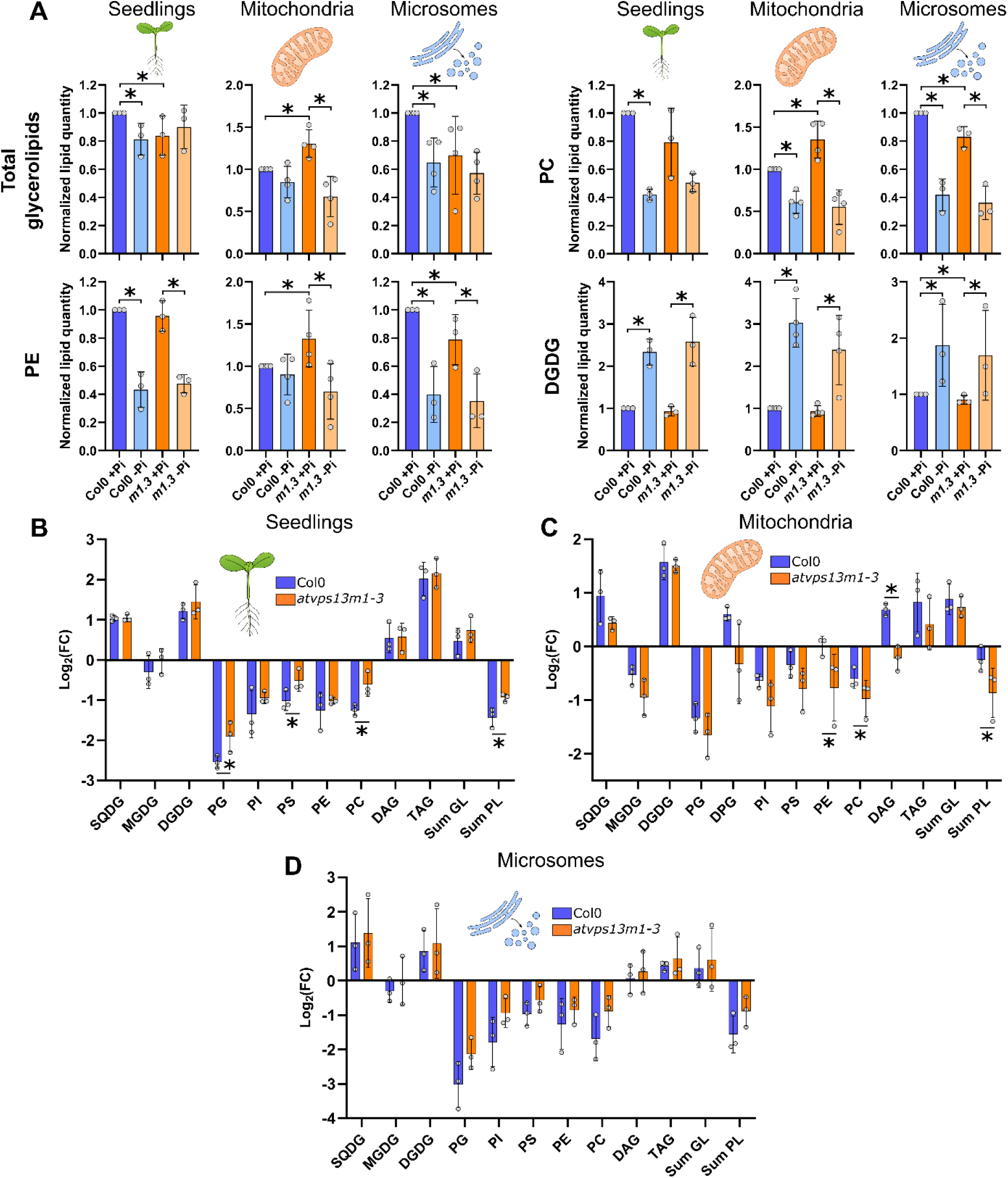
Lipidomic analysis of wild type (Col0) and *atvps13m1-3* seedlings, mitochondria and microsomes isolated from seedlings grown in presence (+Pi) or absence (-Pi) of phosphate (Pi). **A.** Comparison of the normalized lipid quantity for total glycerolipids or isolated lipid classes from Col0 and *atvps13m1-3* mutant seedlings, mitochondria and microsomes obtained in +Pi or -Pi culture conditions. For seedlings, lipids were quantified in nmol per mg of fresh weight while for mitochondrial and microsomal, lipids were quantified in nmol per mg of protein content (see Material and Methods). **B-D**. Log_2_ fold change of each lipid class quantity obtained from seedlings (B), mitochondria (C) and microsomes (D) in -Pi versus +Pi in Col0 and *atvps13m1-3* mutant. On each graph, the means, standard deviations and individual measures from each biological replicate are represented. Statistical analyses were performed using Mann-Whitney tests with a p-value ≤ 0,05 (*).

Lipid class analysis revealed further imbalances. In *atvps13m1-3* microsomes, almost all lipid classes were reduced under Pi-replete conditions, whereas major phospholipids (PC, PE, PI, PG, DPG) accumulated in mitochondria (Fig. 8A, S8), suggesting that intracellular lipid distribution is disrupted. This imbalance was not observed under Pi starvation (Fig. 8A, S8).

To assess lipid remodeling efficiency, we analyzed the Log2FC of each lipid class between 0 mM and 1 mM Pi in all three fractions. As previously observed, phospholipid degradation in whole seedlings was impaired in the mutant, with a similar (though non-significant) trend in microsomes (Fig. 8B, D). In mitochondria, however, phospholipid degradation was more extensive in *atvps13m1-3* than in Col0. This was particularly striking for PE, which was largely preserved in Col0 mitochondria but significantly degraded in *atvps13m1-3* under Pi starvation (Fig. 8A, C). DAG levels provided further insights. Under Pi starvation, DAG increased in whole seedlings of both lines and in Col0 mitochondria, suggesting phospholipid polar head removal at the mitochondrial surface (Fig. 8B, C, Fig. S8). In *atvps13m1-3* mitochondria, however, DAG levels tended to decrease under Pi starvation, pointing to a key role for AtVPS13M1 in coordinating mitochondrial lipid degradation. Additionally, DGDG content remained unchanged in *atvps13m1-3* across all fractions, indicating that AtVPS13M1 does not regulate DGDG level and distribution, which is consistent with its incapacity to bind DGDG *in vitro* (see above).

Overall, our lipidomic analyses reveal that under Pi-replete conditions, the absence of AtVPS13M1 triggers phospholipid accumulation in mitochondria at the expense of microsomes. Under Pi starvation, phospholipid degradation slows in microsomes but accelerates in mitochondria. These results suggest that AtVPS13M1 regulates phospholipid distribution between mitochondria and other organelles.

## Discussion

Our study demonstrates the role of AtVPS13M1 as a lipid transfer protein involved in the regulation of lipid homeostasis, in particular in lipid remodeling in response to Pi starvation in *A. thaliana*. From our data, we conclude that AtVPS13M1 regulates the distribution of lipids between mitochondria and other organelles in both high and low Pi conditions (Fig. 9).

**Figure 9.**
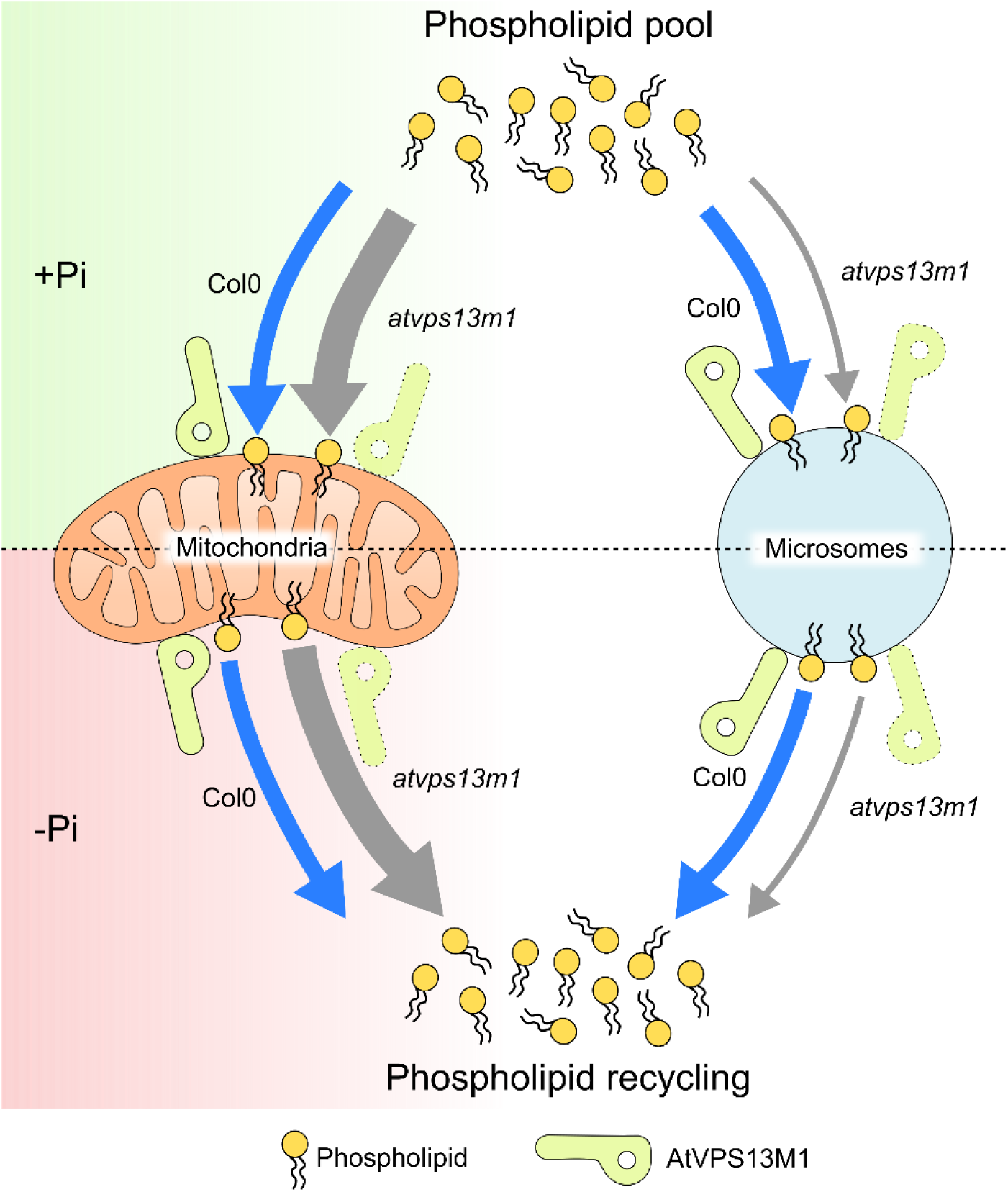
Model of AtVPS13M1 roles in lipid homeostasis in phosphate-replete condition and during lipid remodeling triggered by phosphate starvation in plants. AtVPS13M1 is located at the surface of mitochondria and associates with the microsomal membrane fraction. In phosphate-replete conditions (+Pi), the absence of AtVPS13M1 (*atvps13m1*) causes an accumulation of phospholipids in mitochondria at the expense of the microsomal fraction representing most endosomal compartments. In low phosphate conditions (-Pi), removing AtVPS13M1 promotes mitochondrial phospholipid degradation while phospholipids tend to accumulate in the microsomal fraction. Altogether, our model suggests a general role of AtVPS13M1 in the lipid homeostasis of *A. thaliana* in both phosphate replete and deplete conditions by balancing the lipid supply between mitochondria and other endomembranous compartments.

In calli, AtVPS13M1 is transcribed into one main splice variant, named A, which is not responsive to Pi starvation. Alternative splicing is a major process regulating transcriptomes and proteomes in Eukaryotes. In *A. thaliana*, between 42 and 64% of intron-containing genes produce more than one mRNA ^47,48^, the major alternative splicing event corresponding to intron retention ^48,49^. Our ONT sequencing data revealed the presence of many AtVPS13M1 intron retention-containing alternative spliced variants in *A. thaliana* calli, among which five were confirmed by PCR. The A isoform, also found by ONT RNA sequencing in 14 days-old seedlings ^35^, encodes a full-length protein and is the most abundant form in calli. Most of the other detected isoforms are of very low abundance (< 4%) and produced premature termination codons-containing transcripts. Isoform G retains intron 49, producing a protein containing an additional sequence of 21 aa in the VAB domain. Structural prediction using ColabFold ^50,51^ showed that this extra-sequence does not modify the general folding of the AtVPS13M1 VAB domain into a repeat of 6 modules (Fig. S9A), but increases the size of a loop located close to the C-terminus of the module 4 (Fig. S9B). Therefore, this sequence might regulate AtVPS13M1 localization and/or function by interacting with specific partners. In humans, all the VPS13 genes have at least two major splice variants, but also multiple minor isoforms, some of which contain premature termination codons ^2,8,9,52^. It remains to be determined whether these are splicing errors, likely to occur for long transcripts with numerous introns (63 in AtVPS13M1), or whether they play a key role in the regulation of VPS13 protein levels, localization and/or function.

Redundancy between lipid transport routes is an inherent difficulty in studying LTPs and prevents the appearance of dramatic observable phenotypes. In accordance, *atvps13m1* mutant plants grown in normal conditions presented no obvious growth or root phenotypes, even in low Pi. The absence of AtVPS13M1 did not impair or delay pollen germination neither the degradation of NBR1, suggesting that the protein is not a major player of autophagy in nitrogen and carbon starvation conditions. At present, only a few other studies have conducted phenotypic analyses on VPS13 mutant plants. The *atvps13s* mutant plants have a dramatic phenotype, producing smaller plants with smaller roots and petals, curled leaves, and that are both male and female sterile ^16,17^. This dramatic phenotype of *atvps13s* plants contrasts with the absence of clear developmental phenotype in *atvps13m1* mutants. As AtVPS13M1 and AtVPS13M2 are close paralogs resulting from a gene duplication event during Charophyte evolution and are almost identical in terms of domain organization ^3^, it is therefore very likely that they share common cellular functions and can partially compensate for each other, hence the absence of measurable phenotype in the tested conditions. AtVPS13M2 seems to have acquired specific functions, as mutations in its gene causes a delay in pollen germination and growth, whereas the *atvps13m1* mutants do not display a pollen-associated phenotype ^18^. It is interesting to note that AtVPS13M2 only recently lost the C-terminal β-tripod domain, since this feature is found exclusively in Angiosperms (i.e. flowering plants). Thus, it is possible that AtVPS13M2 retained the general functions of the ancestral protein and acquired new functions during evolution, notably after the loss of the C-terminal β-tripod domain ^3^. Therefore, the generation of an *atvps13m1/atvps13m2* double KO mutant or phenotypic analyses using other stress conditions or developmental stages could help decipher the role of AtVPS13M paralogs in plants in the future.

Yet, despite the absence of macroscopic phenotypes, our work uncovers a critical role for AtVPS13M1 in cellular lipid homeostasis. We show that AtVPS13M1 is not only able to bind a wide range of phospholipids, as already thoroughly described for yeast and human VPS13 proteins ^13,41^, but also neutral lipids (i.e. DAG and TAG), the plastid sulfoglycerolipid SQDG and the galactoglycerolipid MGDG. Hence, our *in vitro* binding experiments suggest that AtVPS13M1 likely transports lipids with a rather low specificity. However, we noticed some peculiarities. First, we did not detect binding to DGDG, suggesting that AtVPS13M1 cannot transport this plastidial galactoglycerolipid. Second, our results suggest that AtVPS13M1 preferentially binds PS species from insect cells, which are enriched in saturated and mono-unsaturated fatty acids, rather than their plant counterparts, which contain more unsaturated acyl-chains. Additionally, we show that AtVPS13M1(1-335) is able to transfer phospholipids *in vitro*, demonstrating the conserved lipid transfer function of VPS13 proteins in plants.

In this work, we show that AtVPS13M1 is primarily located at the mitochondrial surface and also to endomembranes. VPS13 proteins from yeast and humans are known to associate with mitochondria, forming punctate structures or patches on the mitochondrial surface, reflecting their propensity to localize in mitochondrial MCSs with various organelles ^39,41,46,53–55^. Instead of a punctate distribution, we observed a rather uniform AtVPS13M1 coating on the surface of mitochondria, including in actively moving ones. This may suggest that AtVPS13M1 is only transiently engaged in mitochondria-organelle contact sites when it encounters its partners on the surface of other organelles or, alternatively, it may have other MCS-unrelated roles on the mitochondrial surface that remain to be determined. Since AtVPS13M1 harbors a FFAT motif in its N-terminal part, it is tempting to speculate that it is able to interact with VAP-like proteins localized in the ER, such as yeast and human VPS13 proteins ^56^. The localization of VPS13 proteins is a complex process regulated by its numerous functional domains and the expression level of their partners ^1,5^. Given the high number of additional functional domains in AtVPS13M1, this implies that it may have the possibility to associate with various other compartments ^1,3^. While our data makes it clear that AtVPS13M1 associates with mitochondria, much remains to be done to identify the partner organelles and to fully understand what triggers its association with various membranes. Interestingly, AtMIC60, a mitochondrial inner membrane protein located at contact sites with the outer membrane, is involved in the export of phospholipids from mitochondria in response to Pi starvation ^31^. AtMIC60 is part of a larger complex suspected to be found at mitochondria-organelle contact sites to promote lipid transport. Although AtVPS13M1 was not detected in proteomic analysis of the MTL complex ^31^, it may be of future interest to investigate whether AtVPS13M1 could be connected to the MTL complex to support phospholipid trafficking between mitochondrial membranes and other cells compartments.

Recent works have established VPS13 proteins as lipid transfer proteins with pleiotropic roles in lipid-demanding processes ^5,6^. However, loss of VPS13 often causes minimal disruption to whole-cell lipid homeostasis, likely due to redundant transport pathways or context-specific functions in particular cell types, organelles, or stress conditions. For instance, VPS13C has been shown to have an impact on the lysosomal lipidome, whereas lipidome perturbations have only been observed in some brain tissues from patients suffering from Chorea-acanthocytosis, a neurodegenerative disease associated to VPS13A mutation ^57,58^. Similarly, VPS13A or ScVPS13 have been shown to regulate phoshatidylinositol-phosphate levels in the Golgi, plasma membrane or prospore membrane, respectively ^59,60^. Importantly, our lipidomic analyses reveal that AtVPS13M1 regulates phospholipid homeostasis at both whole-cell and subcellular levels. While whole-cell lipidomes show only minor perturbations depending on tissue and developmental stage (Figs. 6, 8, S6, S8), lipid remodeling is consistently impaired in *atvps13m1* mutants across all conditions tested (calli, rosettes, seedlings), and is marked by reduced phospholipid remobilization under low Pi and galactolipid overaccumulation in some tissues.

To pinpoint where this defect occurs, we analyzed mitochondrial and microsomal lipidomes (Figs. 8, S8). Strikingly, loss of AtVPS13M1 has opposite effects on these two fractions. Under Pi-replete conditions, phospholipids accumulate in mitochondria at the expense of microsomes, suggesting AtVPS13M1 normally regulates phospholipid distribution between mitochondria and microsomal membranes. During Pi starvation, however, mitochondrial phospholipids are hyper-remobilized in the mutant, while microsomal phospholipids show reduced degradation compared to wild type, as in the whole-cell. Thus, AtVPS13M1 protects mitochondrial phospholipids from over-degradation during Pi starvation while promoting their turnover in endomembranes (Fig. 9). Yet, the precise mechanisms by which AtVPS13M1 regulates phospholipids distribution remains to be investigated and will be a prospect for future exciting studies. In Pi-repleted condition, AtVPS13M1 could directly regulate phospholipids distribution at ER-organelles contact sites either by transferring lipids, newly synthesized at the ER, to other organelles or by regulating organelle proximity. Alternatively, it could directly exchange phospholipids between mitochondria and other cell compartments. During Pi starvation, AtVPS13M1 regulates phospholipids remodeling in extra-plastidial compartments, favoring the degradation of phospholipids in the endomembranes while preventing the over-recycling of mitochondrial lipids. In contrast, our data showed that AtVPS13M1 does not play a major role in the metabolism and distribution of DGDG during Pi starvation, further supporting the specific role of AtVPS13M1 in phospholipid metabolism and transport rather than galactolipid metabolism.

In conclusion, our data supports a model in which AtVPS13M1 acts as a potent lipid transporter that plays an important role in lipid metabolism and remodeling in response to Pi starvation at the level of mitochondria and extra-plastidial organelles. Further work will be carried out in the future to understand at which other organelles and contact sites AtVPS13M1 operates, how its localization is regulated and what are its specific functions, particularly regarding plant adaptation to Pi starvation.

## Material and Methods

### Plant growth conditions

*A. thaliana* ecotype Col0 was used as wild type and *atvps13m1* T-DNA insertion mutants were obtained from the European Arabidopsis Resource Center NASC: *atvps13m1-1*: SALK_128148 (T-DNA inserted at position 2317, exon 7); *atvps13m1-2*: SALK_019531 (T-DNA inserted at position 4199, exon 14); *atvps13m1-3*: SALK_021158 (T-DNA inserted at position 4985, exon 17); *atvps13m1-4*: SALK_096315 (T-DNA inserted at position 7761, exon 25). T-DNA insertion positions were verified by PCR and sequencing. For plant growth, seeds were surface-sterilized and stratified 3 days at 4°C on dark in water before sowing on soil. Plants are grown on growth chambers under a 16h light, 20°C / 8h dark, 18°C cycle (light intensity 90µmol/m^2^/s, humidity 70%).

### Calli formation, growth and maintenance

*Arabidopsis* seeds were sown and grown on Murashige & Skoog (MS, 4.41 g/L MS (CaissonLab MSP09), 10 g/L sucrose, 0.5 g/L MES 1.2 mg/L, agar 0.8 %, pH 5.7) plates at 22°C under long day conditions. After two weeks of growth, the cotyledons were transferred on MS calli plates (4.41 g/L MS, 30 g/L sucrose, 1.2 mg/L 2,4-dichlorophenoxyacetic acid (2,4D), 0.8 % agar, pH 5.7) and maintained 4 weeks under continuous light at 22°C to obtain calli. Calli were then passed to new MS calli plates once per month. For experiments on liquid media, calli were transferred on 200mL of MS calli liquid media (4.41 g/L MS, 15 g/L sucrose, 1.2 mg/L 2,4D, pH 5.7) and incubated under 125rpm agitation, continuous light at 22°C. Calli were passed in new media every seven days and experiments are started at least six weeks after transfer to liquid media.

### Phosphate starvation on calli

The equivalent of 5mL of sedimented 7 weeks-old calli grown in liquid media were washed 3 times with 50mL of MS +Pi or –Pi liquid media (4.41 g/L MS basal salts -Pi (CaissonLab MSP11), 15 g/L sucrose, 1.2 mg/L 2,4D, Vitamin 1X (Caisson Lab MVL01), +/- 0.5mM Pi (0.6 mM KH_2_PO_4_ + 0.05 mM Na_2_HPO_4_), pH 5.7). Calli are then transfered in 200mL of MS +Pi or –Pi liquid media and incubated under 125rpm agitation, continuous light at 22°C. Around 500 mg or 100 mg of calli were harvested for lipid analysis or RNA extraction, respectively.

### Phosphate starvation on seedlings

Seeds from Col0 and *atvps13m1-3* plants grown and harvested in the same time were surface sterilized with ethanol and bleach and sown on MS medium 1 mM Pi (Duchefa DU1072 1.06 g/L, 1 % (w/v) sucrose, 1 mM KH_2_PO_4_, 1.2% (w/v) agar (Sigma A7921), pH 5.8). Seed were stratified 48h at 4°C on dark and incubated vertically four days in a growth chamber under long day condition (16h light 90 µE / 8h dark) at 22°C. After four days, plants with a similar root length were transferred on MS medium with 5 µM Pi (Duchefa DU1072 1.06 g/L, 1 % (w/v) sucrose, 0.005 mM KH_2_PO_4_, 1.2% (w/v) agar (Sigma A7921), pH 5.8) and placed vertically again in the same growth chamber. At Day 6, rosettes were harvested, fresh weight was measured and rosettes were frozen in liquid nitrogen for lipid analysis.

### Lipid extraction from calli and rosettes

Calli and rosettes were lyophilized overnight using a lyophilizator at -80°C (Christ Alpha 2-4 LD plus). The lyophilized samples were grounded in a glass-tube using a Potter-Elvehjem pestle. 4 mL of boiling ethanol were added to the samples and incubated 5 min at 90°C prior to addition of 2 mL of methanol and 8 mL of ethanol-stabilized chloroform. Samples were homogenized by bubbling argon and incubated 1 h at room temperature before being filtrated through quartz wool. The organic phase containing the lipids is separated from the hydrophilic phase by adding 3 mL of chloroform/methanol (2:1 v/v) and 5 mL of NaCl 1% (w/v). After mixing with bubbled argon, the samples were centrifuged 10 min at 2000 rpm at room temperature and the inferior organic phase is collected. The lipid extracts were retrieved by evaporating the organic phase under argon.

### Lipid extraction from mitochondria and microsomes

Lipids were extracted using a bligh and dyer protocol. Briefly, 1.875 mL of methanol/chloroform 2/1 were added to 300 µg of mitochondria or microsomes in 500 µL of Washing buffer without BSA. After vortexing, 625 µL of H2O and 625 µL of chloroform were added. Samples were vortexed and centrifuged 10 min at 1 000 xg at RT. The inferior organic phase containing the lipids was retrieved and evaporated under argon.

### Methanolysis and total fatty acid methyl esters (FAMEs) quantification by gas chromatography-flame ionization detector (GC-FID)

The lipid extracts are resuspended in 1 mL or 250µL of ethanol-stabilized chloroform for calli, seedlings or rosettes lipids, mitochondria, microsomes, respectively, and 50 µL are mixed with 5 µg of internal standard (C15:0) and 3 mL of 2.5% sulfuric acid in methanol in sealable glass tubes and incubated at 100°C for 1 h to produce FAMEs. Reaction was stopped with 3 mL of water and phase separation was induced by the addition of 3 mL of hexane. After 20min at room temperature, the upper organic phase containing the FAMEs was collected and the phase partition step is iterated. The collected hexane phases were evaporated under argon.

The FAMEs were resuspended in 100 µL of hexane and analyzed by GC-FID (Perkin Elmer Clarus 580) on a 30-m long cyanopropyl polysilphenesiloxane column (SGE BPX70) with a diameter of 0.22 mm and a film thickness of 0.25 μm. The GC column is heated at 180°C and nitrogen is used as vector gas. FAMEs were identified by comparison of their retention times with those of standards (Sigma-Aldrich) and quantified by the surface peak method using the C15:0 fatty acid internal standard for calibration.

### Quantification of glycerolipids by high performance liquid chromatography coupled to triple quadrupole mass spectrometry (HPLC-MS/MS)

The lipid extracts corresponding to 25 nmol of FAMEs were dissolved in 100 µL of chloroform/methanol [2/1, (v/v)] containing 125 pmol of each internal standard. Internal standards used were PE 18:0-18:0 and DAG 18:0-22:6 (Avanti Polar Lipid) and SQDG 16:0-18:0 extracted from spinach thylaKOid (Demé et al., 2014) and hydrogenated as described in ^61^. Lipids were then separated by HPLC and quantified by MS/MS.

The HPLC separation method was adapted from (Rainteau et al., 2012). Lipid classes were separated using an Agilent 1260 Infinity II HPLC system using a 150 mm×3 mm (length × internal diameter) 5 µm diol column (Macherey-Nagel), at 40°C. The mobile phases consisted of hexane/isopropanol/water/ammonium acetate 1M, pH5.3 [625/350/24/1, (v/v/v/v)] (A) and isopropanol/water/ammonium acetate 1M, pH5.3 [850/149/1, (v/v/v)] (B). The injection volume was 20 µL. After 5 min, the percentage of B was increased linearly from 0% to 100% in 30 min and stayed at 100% for 15 min. This elution sequence was followed by a return to 100% A in 5 min and an equilibration for 20 min with 100% A before the next injection, leading to a total runtime of 70 min. The flow rate of the mobile phase was 200 µL/min. The distinct glycerophospholipid classes were eluted successively as a function of the polar head group.

Mass spectrometric analysis was done on a 6470 triple quadrupole mass spectrometer (Agilent) equipped with a Jet stream electrospray ion source under following settings: Drying gas heater: 260°C, Drying gas flow 13 L/min, Sheath gas heater: 300°C, Sheath gas flow: 11L/min, Nebulizer pressure: 25 psi, Capillary voltage: ± 5000 V, Nozzle voltage ± 1000. Nitrogen was used as collision gas. The quadrupoles Q1 and Q3 were operated at widest and unit resolution respectively. PC analysis were carried out in positive ion mode by scanning for precursors of m/z 184 at a collision energy (CE) of 35eV. SQDG analysis was carried out in negative ion mode by scanning for precursors of m/z -225 at a CE of -55eV. PE, PI, PS, PG, PA, MGDG and DGDG measurements were performed in positive ion mode by scanning for neutral losses of 141 Da, 277 Da, 185 Da, 189 Da, 115 Da, 179 Da and 341 Da at CEs of 29 eV, 21 eV, 21 eV, 25 eV, 25 eV, 8 eV and 11 eV, respectively. Quantification was done by multiple reaction monitoring (MRM) with 30 ms dwell time. DAG and TAG species were identified and quantified by MRM as singly charged ions [M+NH_4_]+ at a CE of 19 and 26 eV respectively with 30 ms dwell time. CL species were quantified by MRM as singly charged ions [M-H]- at a CE of -45 eV with 50 ms dwell time. Mass spectra were processed by MassHunter Workstation software (Agilent) for identification and quantification of lipids. Lipid amounts (pmol) were adjusted for response differences between internal standards and endogenous lipids and by comparison with a quality control (QC). QC extract corresponds to a known lipid extract from *Arabidopsis* cell culture qualified and quantified by TLC and GC-FID as described by (Jouhet et al., 2017).

### Extraction of total RNAs from Arabidopsis thaliana calli and RT-PCR

Calli were grounded in liquid nitrogen and approximately 100 mg of powder are collected in 2 mL microcentrifuge tubes before immediately adding 1.5 mL of TRI Reagent® (Sigma-Aldrich). The samples were agitated for 10 min on a rotating wheel and centrifuged at 12 000 x g at 4°C for 10 min. 300 µL of chloroform were added to the supernatant, and samples were left at room temperature for 15 min before a centrifugation at 12 000 x g at 4°C for 10 min. The upper phase was transferred to a fresh 2 mL microcentrifuge tube and 0.4 volumes of isopropanol and 0.4 volumes of high salt solution (1.2 M NaCl, 0.8 M sodium citrate) were added. Samples were gently mixed, incubated for 1h at room temperature to precipitate RNAs and centrifuged at 12 000 x g at 4°C for 15 min. Pellets were washed in 1 mL of cold (4°C) 75% (v/v) ethanol and resuspended in 50 µL of RNAse free (DEPC-treated) water after drying. RNA concentration was measured on a Nanodrop (NanoDrop 2000, Thermo Fisher) and RNA quality was assessed on 0.8% (w/v) agarose gel.

The SuperScript IV VILO reverse transcription kit (Invitrogen) was used to generate cDNAs from the extracted RNAs. Briefly, 1 µg of RNAs or 2µg for qPCR are subjected to a DNAse (ezDNAse) treatment in a final volume of 10µL. Reverse transcription was realized using the SuperScript IV VILO Master Mix that contains degenerate primers in a final volume of 20µL according to manufacturer’s instructions. Reverse transcription occurs by annealing the degenerate primers at 25°C for 1 min, elongation by the reverse transcriptase at 50°C for 10 min and enzyme inactivation at 85°C for 5 min. The final cDNAs have a theoretical concentration of 50 ng/µL and are diluted 10 times before use or for qPCR, 100 ng/µL and were diluted 1/5, 1/25, 1/100 and 1/500.

### qPCR

qPCR was performed on Col0 samples grown 6 days in + or -Pi in a final volume of 10 µL containing 5 µL of PowerSYBR® GreenPCR Master Mix (Applied Biosystem, 4367659), primers (Table S2) at 0.6 µM and 2µL of cDNA on a CFX ConnectTM Real-Time System (BioRad) with the following program: initial denaturation step at 95 ◦C for 10 min and 40 denaturation-amplification-elongation cycles (95 ◦C, 10 s; 60◦C, 10 s; 72 ◦C, 30 s), followed by melting curve assessment (65 ◦C to 95 ◦C, with a 0.5 ◦C increment). Efficiency curves were performed for each primers (AtVPS13M1, GAPDH, MDH) by performing qPCR on four dilutions. AtVPS13M1 and MDH expression levels were normalized by GAPDH. Four biological replicates and three technical replicates were performed for each condition and each primers.

### Nanopore sequencing and analysis

PCR amplification, ONT sequencing data production and analysis were performed at the sequencing platform of the IGFL.

### PCR amplification, library construction and ONT sequencing

cDNAs were prepared from *A. thaliana* Col0 calli cultivated 4 days in presence or absence of Pi, and in duplicate, i.e. four samples in total. Different primer pairs were tested to amplify At4g17140/AtVPS13M1 gene. A primer pair encompassing the region of interest to generate a fragment around 7.65kb was finally selected for further analysis (IsoA_2485F: 5’TTA-GGC-TTT-CAC-TTC-TCA-CC 3’; IsoA_10136R: 5’ CGA-ATC-TTC-ACC-AAA-ACC-AG 3’). The partial AtVPS13M1 gene was amplified by PCR in 45 cycles using LongAmp Hot Start Taq 2X Master Mix (New England Biolabs), from 30/40ng of cDNA per sample in reaction volume of 100µl. PCR conditions were as follows: denaturation at 94°C for 5 min, followed by 45 cycles of 45 sec 94°C, 45 sec 60°C and 8 min at 65°C, and a final elongation of 8 min at 65°C. PCR products were separated on agarose gel and bands of the expected size were cut on the gel using a razor blade (Figure S1A). Nucleospin Gel and PCR clean-up kit (Macherey-Nagel) was employed to purify DNA from the gel. After quantification with Qubit, Oxford Nanopore Technology (ONT) barcoded libraries were constructed according to the ONT amplicon ligation protocol with the SQK-LSK109 kit, and pooled equimolarly before being deposited on a flowcell R9.4.1 (FLO-MIN106) for sequencing with Mk1C device. Fastq reads were generated by MinKNOW 21.05.20 using guppy version 5.0.13 for basecalling. Only pass reads (>Q7) were used for the analysis.

### Data cleaning

Cutadapt (Martin, 2011) was used to filter reads according to primers and length. The program was setup to retain only reads with the expected PCR primers (IsoA_2485F, IsoA_10136R) at both ends and reads between 5kb and 9kb in length to avoid chimeras (--minimum-length 5000 and –maximum-length 9000). In this way, around 50k to 70k reads were retained by sample (**Figure 1B**).

### Identifying isoforms with FLAIR

The *Arabidopsis thaliana* genome (TAIR10) and annotation (TAIR10.49.gtf) were used as a reference to identify isoforms from all reads using the FLAIR pipeline (Tang et al., 2020) and up to the Quantify step. This first analysis identified a lack of one base (T) in the genomic sequence of the TAIR10 reference compared with the long reads generated, and a probably erroneous annotation for at least one exon boundary. As the latter point was also observed in another publication (Zhang et al 2020), we decided to modify the reference sequence and annotation file accordingly. TALON was used to generate the new gtf file (https://github.com/mortazavilab/TALON; Wyman et al., 2019), as FLAIR was unable to identify isoforms from exons not present in the original gtf file. In brief, 1000 reads were randomly sampled from replicate 1 (+Pi) with the seqtk tool (https://github.com/lh3/seqtk) to reduce computation time, the Python program TranscriptClean (https://github.com/mortazavilab/TranscriptClean) (Wyman and Mortazavi, 2019) was first applied to correct mismatches and microindels in long reads mapped to the TAIR genome. TALON was then used with default options on the corrected reads to generate the new gtf file. The FLAIR pipeline was run for a second analysis (module parameters: “align” -v1.3; “collapse” --stringent; “correct” and “quantify” using default) on the four samples and with the new genomic and gtf references. A threshold of 1% of total reads per sample was arbitrary set to retain the main isoforms described by the program, reducing the dataset to 11 isoforms of interest (labelled A to K; **Figure S1B**, and supplementary **Dataset 1**).

### Confirming isoforms with BLAST

To confirm the reality of these FLAIR-predicted isoforms, an independent verification was carried out using a BLAST-based strategy. This comparative sequence approach searched reads for the presence of unspliced introns or the absence of an exon. When an intron was described as present in an isoform but not in isoform A, the sequence of this intron was searched in all reads by BLAST (Altschul et al., 1990) with default parameters. In a similar way, when an exon was described missing in an isoform compared with isoform A, this time the sequence of the 3’ terminal of the previous exon and the sequence of the 5’ terminal of the following exon were concatenated. This concatenated sequence was searched in all read by BLAST with default parameters. The observation of blast hits in these two types of analyses was considered as an indication of the real presence of the intron or the real absence of the exon.

### Experimental validation of isoform by RT-PCR

To validate the presence of unspliced intron in isoform D, E, G, I and J, a PCR was performed on A. thaliana cDNA +Pi using oligos AtMM469-470 and Platinium SuperFi II Master Mix (ThermoScientific). PCR condition was : denaturation 5 min at 98°C, followed by 35 cycles of 10sec 98°C, 30sec 70°C and 8min (+5sec each cycles) at 72°C, and a final elongation of 5 min at 72°C. PCR product at 7,5kb was purified on gel and validation was performed with Phusion (Thermoscientific) and the following oligos : Isoform D: PCR1: AtMM619-620, PCR2: AtMM619-689, Isoform E: PCR1: AtMM621-622, PCR2: AtMM621-690, Isoform G: PCR1: AtMM623-624, PCR2: AtMM623-691, Isoform I: PCR1: AtMM625-626, PCR2: AtMM625-692 and Isoform J: PCR1: AtMM627-628, PCR2: AtMM627-688. PCR condition was : denaturation 5 min at 98°C, followed by 40 cycles of 10sec 98°C, 20sec 70°C and 30sec at 72°C, and a final elongation of 5 min at 72°C. PCR were analyzed on 2% agarose gel.

### Analysis of AtVPS13M1 protein level by MS-based targeted proteomic

Col0 and *atvps13m1* KD calli were grounded in liquid nitrogen and around 50mg of powder were resuspended by pipetting in 250µL of solubilization buffer (50mM Tris-HCl pH8, 2mM EGTA, 100mM NaCl, 1mM dithiothreitol, 5% (p/v) glycerol, 1% (V/V) Triton X100, 1mM phenylmethylsulfoxide, Complete EDTA-Free protease inhibitor tablet (Roche). Samples were incubated 10 min on a rotor wheel at 4°C and centrifuged 5 min at 5 000 x g at 4°C. The supernatant was harvested and protein concentration was determined using Detergent compatible Bradford reagent (Pierce).

Two independent experiments were conducted, the first one analysing WT, *atvps13m1-1* and *atvps13m1-3* lines and the other one analysing WT, *atvps13m1-2* and *atvps13m1-4* lines. Five µg of proteins from total plant extracts were solubilized in Laemmli buffer and stacked in the top of a 4-12% NuPAGE gel (Invitrogen). After staining with R-250 Coomassie Blue (Biorad), proteins were digested in-gel using trypsin (modified, sequencing purity, Promega) as previously described (Casabona et al., 2013). The resulting peptides were analyzed by online nanoliquid chromatography coupled to MS/MS (Ultimate 3000 RSLCnano and Q-Exactive HF, Thermo Fisher Scientific). For this purpose, the peptides were sampled on a precolumn (300 μm x 5 mm PepMap C18, Thermo Scientific) and separated in a 200 cm µPAC column (PharmaFluidics) using a 240 min gradient. The MS instrument was operating in a scheduled targeted mode. The targeted acquisition method combined two scan events corresponding to a full scan event and a time-scheduled Parallel Reaction Monitoring event targeting the precursor ions selected: peptides specific to AT4G17140.1, peptides specific to fructokinases and peptides specific to isoforms of the 26S proteasome regulatory protein subunit RPN5 (Sup. Table 1). The MS and MS/MS data were acquired by Xcalibur version 2.9 (Thermo Fisher Scientific). The mass spectrometry proteomics data have been deposited to the ProteomeXchange Consortium via the PRIDE (Perez-Riverol et al., 2022) partner repository with the dataset identifier PXD052019.

Peptides and proteins were identified by Mascot (version 2.8.0, Matrix Science) through concomitant searches against the TAIR database (version 10.0) and a homemade database containing the sequences of classical contaminant proteins found in proteomic analyses (keratins, trypsin, etc.). Trypsin/P was chosen as the enzyme and two missed cleavages were allowed. Precursor and fragment mass error tolerances were set at respectively at 10 and 20 ppm. Peptide modifications allowed during the search were: Carbamidomethyl (C, fixed), Acetyl (Protein N-term, variable) and Oxidation (M, variable). The Proline software (Bouyssié et al., 2020) was used for the compilation, grouping and filtering of the results (conservation of rank 1 peptides, peptide length ≥ 6 amino acids, false discovery rate of peptide-spectrum-match identifications < 1% (Couté et al., 2020), unique interpretation of spectrum, and minimum of one specific peptide per identified protein group). Proline was then used to perform a MS1-based label-free quantification of the identified peptides.

### Recombinant protein expression and purification from insect cells

The AtVPS13M1(1-335)-6xHis fragment was cloned in the pACEBac1 vector (Geneva Biotech) by Gibson assembly. One µg of vector was introduced by heat shock in *E. coli* DH10EMBacY competent cells carrying the baculovirus genome (bacmid) and transformant were selected on LB supplemented with 50 µg/mL kanamycin, 10 µg/mL tetracyclin, 10 µg/mL gentamicin, 1 mM IPTG and BluoGal 200 µg/mL. White colonies, in which the gene of interest was transferred to the baculovirus genome, were streak out on fresh selection plate. Plasmid was extracted from an overnight 2 mL culture and precipitated with isopropanol. After washing with ethanol 70% (v/v), the plasmid was resuspended in sterile water and mixed with Sf21 culture media (ref) and X-tremeGENE HP DNA transfection reagent (Roche). The mix was next added to Sf21 insect cells culture at 0.5x10^6^ cells/mL for transfection. The supernatant containing the recombinant baculovirus (V_0_) was harvested after incubating the cells 72h at 27°C under agitation and used to inoculate fresh Sf21 cells at 0.5x10^6^ cells/mL (3 mL of V_0_ for 25mL of cells). Infection was followed daily by cell counting and YFP observation. After 72h, cells have stopped dividing and supernatant (V1), containing high amount of recombinant baculovirus, was harvested and stored at 4°C. For large scale production, Sf21 cells at 0.5x10^6^ cells/mL were inoculated with V1 (1/1000) and incubated at 27°C under agitation. Cells were harvested after 5 days and pellet were stored at -80°C.

For protein purification, cells were resuspended in Lysis buffer (50 mM Tris-HCl pH7.5, 100 mM NaCl, 10 mM MgCl_2_, 10 mM imidazole, 1 mM PMSF and protease inhibitor cocktail EDTA-free (Roche) and sonicated 15 min (output 50%, duty cycle 7) on ice. After a centrifugation 20 min at 40 000 x g 4°C, the protein was mostly insoluble and in the pellet fraction. The pellet was gently resuspended in Lysis buffer + 0.2% (w/v) Brij35 and incubated 20 min on ice. After a centrifugation of 10 min at 40 000 x g 4°C, the supernatant, containing the recombinant protein, was harvested and loaded on Ni Sepharose 6 Fast Flow (Cytiva) pre-equilibrated with Lysis buffer + 0.2% (w/v) Brij35. The column was washed with Washing buffer 1 (50 mM Tris-HCl pH7.5, 100 mM NaCl, 10 mM MgCl_2_, 25 mM imidazole, Brij 35 0.02% (w/v)) and Washing buffer 2 (50 mM Tris-HCl pH7.5, 100 mM NaCl, 10 mM MgCl_2_, 65 mM imidazole, Brij 35 0.02% (w/v)) before protein elution with Elution buffer (50 mM Tris-HCl pH7.5, 100 mM NaCl, 10 mM MgCl_2_, 300 mM imidazole, Brij 35 0.02% (w/v)). After concentration on 30K Amicon Ultra Centrifugal filter (Millipore), 500 µL of protein were loaded on a Superose 6 increase 10/300 GL (Cytiva) size exclusion column pre-equilibrated with SEC buffer (50 mM Tris-HCl pH7.5, 150 mM NaCl, 10 mM MgCl_2_, Brij 35 0.02% (w/v)) and using an AKTA Pure^TM^ chromatography system. Separation was performed at 4°C with a flow rate of 0.5 mL/min and 1mL fractions were harvested. Fractions containing recombinant protein were pooled and concentrated on a 30K Amicon Ultra Centrifugal filter (Millipore). Glycerol at 10% final was added to the sample and protein concentration was estimated using Bradford detergent compatible assay (Pierce). The protein was used fresh for transfer assays or was stored at -80°C for lipid binding assays. Tom20.3 soluble domain was purified as described in (Michaud et al., 2016).

### Fluorescent in vitro lipid binding assay

20 µg of purified proteins were incubated with 1 µg of commercial nitrobenzoxadiazole (NBD)-conjugated lipids in lipid binding buffer (50 mM Tris pH 8, 300 mM NaCl, 10% (v/v) glycerol) at 22°C for 1h. The NBD-conjugated lipids were NBD-PC, NBD-PE, NBD-PS and NBD-PA (Avanti polar lipids). The samples were run on a commercial CN-PAGE gel (Mini-Protean TGX, native gel, Bio-Rad) at 90V until bromophenol blue reached the bottom of the gel. NBD fluorescence was read directly in-gel using a fluorescence imager (Amersham Typhoon) using a 488 nm excitation laser and a Cy2 emission filter. Protein content was read by Coomassie blue staining (InstantBlue® Coomassie Protein Stain (ISB1L), ABCAM).

### In vitro lipid binding assay

Equivalent of 100 nmol of total lipids extracted from Col0 calli grown in absence of Pi was resuspended in 20 µL of methanol and 800µL of SEC bufer was then added to solubilized lipids. After vortexing, 4.8 nmol of purified recombinant protein in 200 µL of SEC buffer were added to obtain a final volume of 1 mL. After a 60 min incubation at room temperature on a rotor wheel, 200 µL of NiNTA resin were added followed by another 60 min at room temperature on a rotor wheel. The resin was washed two times with 1 mL of SEC buffer and protein elution was performed with 550 µL of SEC buffer with 300 mM imidazole. 500 µL of sample were used for lipid extraction and LC-MS/MS analysis and the rest was used to evaluate protein concentration using Bradford a ssay. For lipid extraction, 1.875 mL of CHCl_3_/MeOH 1/2 were added on the samples from *in vitro* lipid binding assays (500 µL). After vortexing, 625 µL H_2_O and 625 µL CHCl_3_ were added and samples were vortexed and incubated 10 min at room temperature. After a centrifugation of 10 min at 1000 x g, the organic phase was retrieved and transferred in a new tube and dried under argon. Lipid films were stored at -20°C.

### In vitro fluorescent lipid transfer assay

In vitro fluorescent lipid transfer assays were adapted from (Kumar et al. 2018). Lipids to form donor (61% POPC, 30% soy PE, 2% NBD-PS, 2% rhodamine-PE and 5% DGS-NTA Ni) or acceptor (70% POPC, 30% soy PE) liposomes were mixed and dried under argon. Lipid films were resuspended in liposome buffer (Hepes-KOH 50 mM pH 7,4, NaCl 100 mM) at 1mM and 6mM for donor and acceptor liposomes, respectively. After hydration by incubating 1h at RT with regular vortexing, lipids were submitted to 5 cycles of freeze/thaw in liquid nitrogen and 50°C water bath. Liposomes were extruded through a 100nm polycarbonate membrane using a mini-extruder (Avanti Polar Lipid). Liposomes were kept on ice and used within 48h for i*n vitro* lipid transfer assays.

*In vitro* transfer assays were performed in 200 µL with 25 µM of donor liposomes, 0 to 1.5 mM of acceptor liposomes in liposome buffer. Reaction started by the addition of the purified recombinant protein at a final concentration of 0,125 µM. Assays were performed on black microplates on a TECAN SPARKS M10 with an excitation at 460 nm and emission at 538 nm. Fluorescence signal was measured every 60 sec during one hour with a 3 sec agitation at 56 rpm 3 sec before reading. Fluorescence values were normalized to the value obtained at T=0.

### AtVPS13M1 tagging by recombination

Recombination-based 3xYepet and GUS tagging of AtVPS13M1 was performed using the strategy and protocols described in (Alonso and Stepanova, 2014; Brumos et al., 2020). Transformable Artificial Chromosome N°78L17 from the JaTY collection (Hirose et al., 2015) containing the At4g17140 gene was obtained from the Arabidopsis Biological Resource Center (ABRC, https://abrc.osu.edu/) and the presence of the gene was verified by PCR. JAtY78L17 contains a gDNA fragment of 70Kb with 14Kb and 32Kb upstream and downstream of the AtVPS13M1 coding sequence, respectively. Tags were introduced between nucleotides 17,557 and 17,558 of the JAtY78L17 gDNA fragment corresponding to the tag insertion after the aa 472 on AtVPS13M1 protein sequence. Plasmid was introduced in E. coli SW105 strains, kindly provided by the Frederick National Laboratory for Cancer Research (Frederick, Maryland, USA), by electroporation as described in (Alonso and Stepanova, 2015) and positive clones were selected by PCR using oligo AtMM631-634. Cassettes containing the 3xYepet or GUS tag were amplified using oligonucleotides containing 40 nt flanking sequences corresponding to the sequence upstream nt 17557 for the forward oligo (AtMM724) or downstream nt 17558 for the reverse oligo (AtMM755). Vectors containing the 3xYepet (JMA690, CD3-1727) or GUS (JMA2352, CD3-2823) cassettes described in (Brumos et al., 2020) were obtained from the ABRC. PCR products were purified and introduced into the SW105-JAtY78L17 strain by electroporation after an incubation of the strain 15 min at 42°C to induce expression of recombination enzymes (Alonso and Stepanova, 2015). Recombined clones were selected on selective media containing carbenicillin and kanamycin at 28°C for two days and confirmed by PCR using oligo AtMM473-726. Excision of the selection Ampicilin marker was induced by the addition of Arabinose which triggers expression of flippases (Alonso and Stepanova, 2015). After selection on kanamycin plates, excision of the selection marker was confirmed by PCR using oligos AtMM473-726. Plasmids were then purified and sequence from the AtVPS13M1 start to stop codon was verified by Sanger sequencing. For the AtVPS13M1-3xYepet construct, plasmid was trimmed in order to reduce plasmid size (Brumos et al., 2020). FRT5-Amp-FRT5 trimming cassette (JMA2372, CD3-2826) was amplified using oligo AtMM737-738 with flanking region allowing insertion between the plasmid and the nucleotide 4 214 of the gDNA. Cassette was inserted in the JaTY78L17-AtVPS13M1-3xYepet plasmids by electroporation as described previously. Recombined clones were selected on kanamycin and carbenicilin plates and controlled by PCR using oligos AtMM723-730. Then, FRT2-Tet-FRT2 trimming cassette (JMA1317, CD3-2827) was amplified using oligo AtMM720-740 with flanking region allowing insertion between the nt 44 289 of the gDNA and the plasmid. Cassette was inserted in the JaTY78L17-AtVPS13M1-3xYepet plasmid containing the first trimming cassette by electroporation as described previously. Recombined clones were selected on kanamycin, carbanecilin and tetracyclin plates and controlled by PCR using oligonucleotides AtMM721-728. Both cassettes were then removed by the induction of flippases expression using L-arabinose and colony selection on kanamycin plates. Excision of the Amp and Tet cassettes were controlled by PCR using oligonucleotides AtMM723-730 and AtMM728-721. Thereafter, JaTY78L17-AtVPS13M1-3xYepet (trimmed) and JaTY78L17-AtVPS13M1-GUS plasmids were introduced by electroporation in *Agrobacterium tumefaciens* strain GV3101 (pMP90) (Farrand et al., 1989) and transformed in *atvps13m1-3* mutant background by floral dip (Clough and Bent, 1998; Alonso and Stepanova, 2014). Basta-resistant plants were selected on soil and confirmed by PCR using oligos AtMM473-726. Plants expressing the constructions were screened by confocal imaging or GUS staining and brought to the F3 used for experiments.

### GUS staining

For analysis in liquid media and Pi starvation condition, seeds were sterilized with ethanol and bleach and washed with sterile water. Around 20 seeds were added in six-wells plates filled with 5mL of liquid media MS+Pi (4.23 g/L MS media (Duchefa DU1072), 10 g/L sucrose, 1mM KH_2_PO_4_, pH 5.8). Seeds were germinated and incubated under continuous light at 22°C and 100rpm for 3 days. A first batch of seedlings was harvested for GUS staining, a second batch was kept in MS+Pi media and a third batch was washed with water and MS-Pi media (4.23 g/L MS media (Duchefa DU1072), 10 g/L sucrose, pH 5.8) two times before transfer to a six-well plate containing 5mL of MS-Pi liquid media. Seedlings in MS+Pi and MS-Pi were further incubated under continuous light at 22°C and 100rpm and samples were harvested for staining after 3 and 6 days of incubation. For growth on vertical plates, after sterilization, seeds were sow on MS+Pi media containing 0,8% (w/v) agar (Sigma A7921) and stratified 3 days at 4°C on dark. Plates were placed vertically in the growth chamber under a 16h light/8h dark cycle at 22°C/20°C and harvested for staining after 7 days. For plant tissues staining, different plant tissues were harvested after 7 weeks of growth on soil for staining the same day in parallel.

Seedlings or plant tissues were harvested and immediately immersed in cold acetone 90% (v/v) for 20 min and then washed once with Staining buffer (50 mM sodium phosphate buffer pH7.4, Triton X100 0.2% (v/v), 2 mM C_6_N_6_FeK_3_ 2 mM C_6_N_6_FeK_4_) and immersed in Staining buffer + 5 mM of X-gluc (Thermoscientific R0852). Samples were vacuum-infiltrated 3 times 10 min at 37°C and samples were then incubated 8 or 24h at 37°C for seedlings and 7 weeks-old plant tissues, respectively. Tissues were washed twice with water and with two baths of ethanol 70% (v/v). Images were acquired using a Keyence VHX-5000 microscope with a VH-Z100R objective.

### Confocal imaging

F3 plants from AtVPS13-3xYepet transformant lines n°6 and 10 were grown on soil in a 16h/8h light/dark cycle and imaged at 7 weeks. For mitochondria staining, plant tissues were harvested and immerged in MitoTracker^TM^ Red FM (ThermoFisher Scientific, M22425) 500 nM in MS media and infiltrated under vacuum 3 times 5 min and further incubated for 2h before washing with water and confocal observation. Laser scanning confocal microscopy was performed on a microscope Zeiss LSM880 with a 63x/1.4 oil-immersed Plan-Apochromat objective, running Zen 2.3 SP1 acquisition software (Platform μLife, IRIG, LPCV). AtVPS13M1-3xYepet was exiting at 514 nm at 25% laser capacity and fluorescence was detected at 520-550 nm with a photomultiplier tube detector set to 800V. MitoTracker^TM^ Red FM was exiting at 561 nm at 1-4% laser capacity and fluorescence was detected at 565-620 nm with a photomultiplier tube detector set to 800V. Images were visualized and treated with the ImageJ 1.53a software running the image processing package Fiji.

### Subfractionation of seedlings grown in liquid media

Col0 and *vps13m1-3* seeds were surface sterilized 2 min in ethanol 70% followed by 15 min in bleach 4% + Tween 20 4% (p/v) under a rotating wheel. After four washes in sterile water, seeds were transferred in MS-Mito +Pi media (Duchefa DU1072 2.1 g/L, 2.5 % (w/v) sucrose, 1 mM KH_2_PO_4_, 0.5 g/L MES, pH 5.8) and grown 4 days at 22°C under 125 rpm agitation and continuous light (50µE). Seedlings were washed three times with MS-Mito +Pi or MS-Mito -Pi media (Duchefa DU1072 2.1 g/L, 2.5 % (w/v) sucrose, 0.5 g/L MES, pH 5.8) and grown again for three days in the corresponding media in the same condition described above. Seedlings were filtered. A sample was retrieved for total lipid analysis and heated at 95°C for 10 min to inactivate phospholipases. Seedling were ground in a cold mortar in Grinding buffer (0.45 M sucrose, 25 mM Na_4_P_2_O_7_, 2 mM EDTA, 10 mM KH_2_PO_4_, 1 % (w/v) PVP-40, 1 % (w/v) BSA, 20 mM sodium ascorbate, 20 mM cysteine, pH 7.5) and filtered into a 50 µM nylon mesh. Samples were centrifuged at 1600 xg for 5 min at 4°C. The supernatants were transferred in clean tubes and were centrifuged at 2500 xg for 5 min at 4°C followed by a centrifugation at 17 200 xg for 20 min at 4°C. For mitochondria purification, pellets were gently resuspended in Washing buffer (0.3 M sucrose, 10 mM MOPS, 0.1 % (w/v) BSA, pH 7.5) and loaded at the surface of PVP-40 0-4.4 % (w/v) continuous gradients (0.3 M sucrose, 10 mM MOPS, 28% (v/v) Percoll, PVP-40 0-4.4% (w/v) pH 7.5). Gradients were centrifuged at 40 000 xg 40 min at 4°C and the mitochondria band was harvested in the lower heavy part of the gradient. Mitochondria were washed three times with Washing buffer without BSA at 31 000 xg 15 min at 4°C and resuspended in Washing buffer without BSA. For microsome preparation, the supernatant obtained after pelleting crude mitochondria at 17 200 xg for 20 min at 4°C was further centrifuged 1h at 100 000 xg at 4°C and membrane pellets obtained resuspended in Washing buffer without BSA. Protein concentration of each fraction was determined using Bradford Reagent assay (BioRad). For lipid extraction, 300 µg of protein was heated 10 min at 95°C to inactivate phospholipases before extraction.

### Subfractionation of calli cell culture, thermolysin treatment and western blot

Mitochondria were extracted from calli grown 4 days in MS+Pi according to a protocol described in ^62^. Briefly calli were grounded in Grinding buffer (0.3 M mannitol, 15 mM 3-(N-morpholino)propanesulfonic acid (MOPS) pH 8, 2 mM ethylene glycol-bis(2-aminoethylether)-N,N,N’,N’-tetraacetic acid (EGTA) pH 8, 0.6 % (w/v) polyvinylpyrrolidone K25 (PVP25), 0.5 % (w/v) bovine serum albumin (BSA), 1 mM dithiothreitol (DTT), 1 mM phenylmethylsulfonyl fluoride (PMSF), 5 mM aminocaproic acid, 1 mM benzamidine) and the mixture was centrifuged at 700 xg for 5 min at 4°C. The supernatant was centrifuged twice 3 000 xg for 5 min at 4°C and the enriched mitochondria fraction was pelleted at 20 000 xg for 15 min at 4°C. The surpernatant was centrifuged at 100 000 xg for 1h at 4°C to separate the soluble fraction (supernatant) from the pelleted microsomes. The mitochondria pellet was resuspended in Washing buffer (0.3 M mannitol, 10 mM MOPS pH 7.4, 1 mM PMSF, 5 mM aminocaproic acid, 1 mM benzamidine), loaded on top of a discontinuous Percoll gradient (5 mL 40%, 20 mL 23%, 10 mL 18%) and centrifuged at 50 000 xg 45 min at 4°C. Mitochondria were harvested at the 40/23% interface and washed three times with Washing buffer. Protein concentration of each fraction was determined using Bradford Reagent assay (BioRad).

For thermolysin treatment, mitochondria were washed in Washing buffer without protease inhibitors (0.3 M mannitol, MOPS 10 mM, pH 7.4). 50 µg of proteins were incubated 1h on ice with 0 to 5 µg of thermolysin (Sigma Aldrich) in washing buffer without protease inhibitors containing 1 mM of CaCl_2_ in a final volume of 200 µL. After incubation, 75µL of NuPAGE LDS 4X (ThermoFisher) and 25 µL of NuPAGE Reducing 10X (ThermoFisher) were added.

For sub-cellular localization analyses, 20 µg of Total calli extract and 10 µg of Mitochondria, Microsomes and Soluble fractions were loaded on gel. For thermolysin treatments, 48 µL of sample were loaded (8.5 µg of proteins). All the samples were loaded on a 4-12% Bolt Bis-Tris Plus NuPAGE gels (ThermoFisher) and migrated in 1X MES-NuPage running buffer (ThermoFisher). Gels were transferred on nitrocellulose membranes for 1h at 25V in NuPAGE Transfer buffer (Thermofisher). Western blots were performed on membranes blocked in TBS-T Milk 5% (w/v) and all the steps were performed in this buffer except for the final washing step which was performed in TBS-T. The antibodies used are : anti-GFP-HRP (Miltenyi Biotech), anti-CoxII, anti-Kar2, anti-Vph1 and anti-UGPase were purchased from Agrisera, anti-Mic60 was described in ^31^ and anti-Tom20.3 was kindly provided by Monika Murcha (University of Western Australia). Western blots were revealed using Clarity ECL substrates (BioRad) on an ImageQuant800 visualizer (Cytiva Life Science).

### Root architecture analysis

Seeds from Col0, *atvps13m1-2* and *m1-3* mutants were surface sterilized with ethanol and sow on plates (half-strength Murashige and Skoog (MS) medium (Duchefa, Netherlands), including 1% (w/v) sucrose, 2.56 mM MES and 1% (w/v) Phytoagar, pH 5.7). Plates were stratified at 4 °C for 2 days in the dark and grown vertically at 22 °C under a 16 h light/8 h dark-cycle. Root growth was determined after 7 days. Live cell imaging was performed using a Zeiss LSM880, AxioObserver SP7 confocal laser-scanning microscope (INST 248/254-1). Bright field images were acquired with a Plan-Apochromat 20x/0.8 M27 objective and propidium iodide-stained roots with a Zeiss C-Apochromat 40×/1.2 W AutoCorr M27 water-immersion objective. For propidium iodide staining, roots were mounted in 0.01 mg/ml PI solution and captured with 543 nm excitation and 580–718 nm emission wavelength. For processing, either the Zeiss software ZEN 2.3 or the Fiji software (https://imagej.net/Fiji) was used.

### Root growth assessment under low Pi

Seeds from Col0*, atvps13m1-3* and *m1-4* plants grown and harvested in the same time were surface sterilized with ethanol and bleach and sown on MS medium 1 mM Pi (Duchefa DU1072 1.06 g/L, 1 % (w/v) sucrose, 1 mM KH_2_PO_4_, 1.2% (w/v) agar (Sigma A7921), pH 5.8). Seeds were stratified 48h at 4°C in the dark and incubated vertically four days in a growth chamber under long day condition (16h light 90 µE / 8h dark) at 22°C. After four days, plants with a similar root length were transferred on MS medium 1 mM Pi or MS medium with 5 µM Pi (Duchefa DU1072 1.06 g/L, 1 % (w/v) sucrose, 0.005 mM KH_2_PO_4_, 1.2% (w/v) agar (Sigma A7921), pH 5.8) and placed vertically again in the same growth chamber. Plates were scanned at day 0 and then at day 1, 2, 3 and 6 on a GS-800 Calibrated Densitometer (BioRad). Primary root length was measured each day using ImageJ SmartRoot plugging and data was normalized by the size of the root at day 0.

### Autophagy assessment by NBR1 level analysis

Seeds were vernalized in water and darkness, at 4 °C, for 24h. The seeds were surface sterilized in 10% bleach for 30 min, and sown on Murashige and Skoog (MS) agar medium plates (4,4 g.L−1 MS powder including vitamins (Duchefa Biochemie M0222), 0.8% plant agar (Duchefa), 1% sucrose and 2.5 mM 2-(N-morpholino)-ethanesulphonic acid (MES), pH 5.7). Seedlings were grown vertically for 7 days, at 21°C, under long-day conditions (16 h-light/8 h-dark photoperiod, 300 μEm²s−1). To induce autophagy, seedlings were starved of nutrients by transferring them from MS plates into liquid MS −NC medium (Murashige and Skoog liquid medium depleted for nitrogen and carbon: Murashige and Skoog micronutrient salts (Sigma, M0529), 3mM CaCl_2_, 1.5 mM MgSO_4_, 5 mM KCl, 1.25 mM KH_2_PO_4_, 0.5% (w/v) D-mannitol, 3 mM MES, pH 5.7), and incubated in darkness (wrapped in aluminum foil) (-NC). For rich conditions, seedlings were incubated in rich liquid Murashige and Skoog (MS) medium (4,4 g.L−1 MS powder including vitamins, 1% sucrose, and 2.5 mM MES, pH 5.7) under normal light conditions (+N).

To assess NBR1 protein levels, 7-day-old seedlings were transferred from full MS plates into liquid medium (+N) or (−NC) for 6 hours and *atg5* KO was used as a control. After treatments, whole seedlings were frozen in liquid nitrogen, disrupted and homogenized in the following buffer: HEPES 50 mM pH 7.5, Sucrose 0.225 M, MgCl2 2.5 mM, DTT 0.5 mM, PVP 0.25% [w/v], PMSF 1 mM, antiprotease mix (P9599 Sigma). Homogenates were centrifuged at 1600 × g for 20 min at 4 °C and twice at 1600 × g for 10 min at 4°C, transferring only the supernatant at each stage. To analyze samples, the protein concentration of each sample was determined using Bio-Rad Protein Assay Dye Reagent Concentrate, (BioRad) and measuring sample absorption at 595 nm. Equal amounts of proteins were prepared, denatured using Laemmli buffer, loaded onto 12% SDS-PAGE, and analyzed by immunoblot using an anti-NBR1 antibody (Agrisera, 1/5000; upper part of the immunoblot) and an anti-Actin (Agrisera, 1/4000^e^) or an anti-UGPase (Agrisera, 1/10000; bottom part of the immunoblot) and a peroxidase-coupled Goat anti-Rabbit secondary antibody. NBR1, Actin, and UGPAse signals were measured using the Image J 1.8.0_172 software’s (National Institutes of Health, USA, http://imagej.nih.gov/ij). Band intensities were calculated using the plot surface areas functionality of the software. NBR1 levels were quantified using signals from total protein (stain-free) and Actin or UGPase for normalization.

### Statistical treatment of data and graphic representation

Statistical significance of data and histograms were performed using GraphPad software (version 6) with a non-parametric Mann-Whitney test.

## Supplemental material

**Figure S1.** PCR of AtVP13M1 fragment used for Oxford Nanopore Technology (ONT) long-read sequencing and results of the FLAIR analysis performed on replicates 1 and 2 in presence or absence of phosphate.

**Figure S2:** Analysis of AtVPS13M1 expression in *Arabidopsis thaliana atvps13m1-3* mutant line stably expressing AtVPS13M1 fused to β-glucuronidase at position 472 expressed under its native promotor

**Figure S3.** Analysis of lipids bound to AtVPS13M1(1-335) fragment *in vitro*.

**Figure S4.** Analyses of AtVPS13M1 expression in *atvps13m1* T-DNA insertion lines.

**Figure S5:** Phenotype analysis of *atvps13m1* mutant plants.

**Figure S6.** Lipid analysis in wild type (Col0) and *atvps13m1-2* KD calli grown in presence and absence of phosphate (Pi) for 6 days and of Col0 and *atvps13m1-3* KD rosettes harvested after 6 days of growth in 1 mM or 5 µM Pi.

**Figure S7:** AtVPS13M1 subcellular localization in 6 weeks-old leaves from *Arabidopsis thaliana atvps13m1-3* mutant stably expressing AtVPS13M1 fused to 3xYepet at position 472 expressed under its native promotor.

**Figure S8:** Lipid analysis in wild type (Col0) and *atvps13m1-3* mutant seedlings, mitochondria and microsomes isolated from seedlings grown in presence (+Pi) or absence (-Pi) of phosphate for 4 days.

**Figure S9.** Alignment of VAB structural prediction of AtVPS13M1 isoform A (in green) and G (in red).

**Table S1.** Results of MS-based targeted proteomic analyses.

**Table S2.** Primers used in this study

**Dataset 1.** Alignment of the AtVPS13M1 isoforms described in TAIR10, found in Zhang et al. 2020 and identified to AtVPS13M1 gDNA.

## Acknowledgments

We thank Emmanuel Thevenon (LPCV, Grenoble), Claude Alban (LPCV, Grenoble), Sylvaine Roy (LPCV, Grenoble) and Vangeli Geshkovsky (LPCV, Grenoble) for their advices for insect cell protein expression, SEC optimization, lipid data processing and GUS staining, respectively. This work was supported by the French National Research Agency in the framework of the “investissement d’avenir” program (ANR-15-IDEX-02) and of the ANRJCJC MiCoSLiT (ANR-19-C13–0013) to MM. It also received funding from GRAL a program from the Chemistry Biology Health (CBH) Graduate School of University Grenoble Alps (ANR-17-EURE-0003). The LIPANG (Lipid analysis in Grenoble) platform is supported by the Rhône-Alpes Region, the fonds FEDER, and GRAL, financed within the University Grenoble Alpes graduate school (Ecoles Universitaires de Recherche) CBH-EUR-GS (ANR-17-EURE-0003). We thank the microscopy facility MuLife of IRIG/DBSCI, funded by CEA Nanobio and GRAL LabEX (ANR-10-LABX-49-01) financed within the University Grenoble Alpes graduate school CBH-EUR-GS (ANR-17-EURE-0003). The proteomic experiments were partially supported by Agence Nationale de la Recherche under projects ProFI (Proteomics French Infrastructure, ANR-10-INBS-08) and GRAL, a program from the Chemistry Biology Health (CBH) Graduate School of University Grenoble Alpes (ANR-17-EURE-0003).

## Data availability

Most of the data are presented as supplementary files or can be provided upon request.

## Author contributions

MM and SL conceived the strategy, the experiments and analyzed the data. SB and YC performed proteomic analyses. BG, JD and SH performed ONT sequencing. AB and JC performed autophagy tests. DS performed root morphology analysis. JJ, MS and MM performed lipidomic analyses. Other experiments were performed by SL or MM with the technical support of CA. MM supervised the project. MM and SL wrote the manuscript with the input from all the authors. All authors reviewed and approved the manuscript.

