## Supplementary figures for "AtVPS13M1 regulates subcellular lipid distribution and lipid remodeling in phosphate-deprived plants"

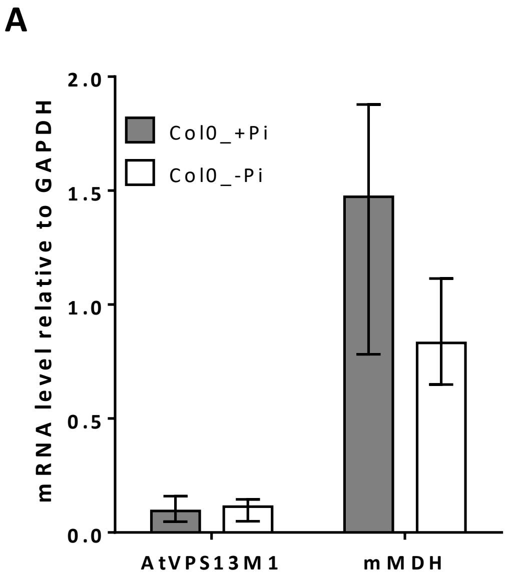


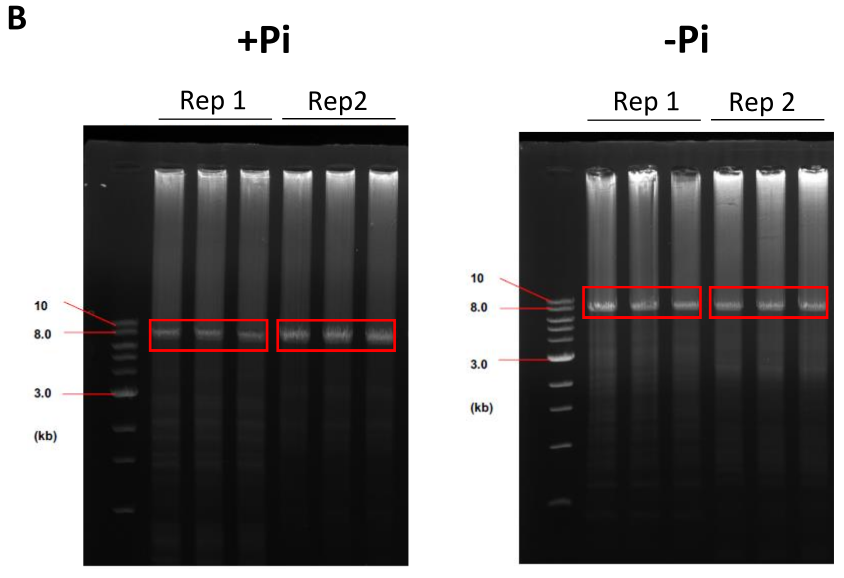


C

| Name | Isoform | Rep 1_+Pi | Rep 1_-Pi | Rep 2_+Pi | Rep 2_+Pi |
|---|---|---|---|---|---|
| TALONT00054017_TALONG000032834 | A | 52 380 | 44 792 | 45 963 | 38 313 |
| 3fbacfbc-998f-4c1b-9229-8657dca25aa0_TALONG000032834 | B | 2 545 | 2 174 | 2 269 | 1 885 |
| 85b91dd2-eb04-4762-8651-62d734afd00d_TALONG000032834 | C | 1 410 | 1 269 | 1 271 | 1 021 |
| 1481f6c7-dcae-4e67-aba5-82575b18fd8e_TALONG000032834 | D | 1 258 | 1 085 | 1 197 | 916 |
| dc0fdf85-f0c3-4b7b-85b0-0c276c3b94d7_TALONG000032834 | E | 965 | 870 | 863 | 750 |
| e507081b-3726-4c22-bed1-81de6b86dbe9_TALONG000032834 | F | 874 | 671 | 764 | 605 |
| 499abbbb-b888-4d3c-a878-eaf14acbcac4_TALONG000032834 | G | 821 | 753 | 686 | 560 |
| 86487668-56b8-41eb-8f3d-67ab19935935_TALONG000032834 | H | 800 | 658 | 728 | 620 |
| d6c2ec2f-420d-43e4-a291-3f2a37e00a17_TALONG000032834 | I | 796 | 723 | 683 | 610 |
| 478bfef4-b6fa-4806-8b91-82e6033240e6_TALONG000032834 | J | 762 | 728 | 660 | 507 |
| cdfac354-c3ee-4ec6-b77e-5cb5072f13c4_TALONG000032834 | K | 719 | 632 | 641 | 574 |

**Figure S1. Analysis of AtVPS13M1 transcripts in response to phosphate (Pi) starvation in *Arabidopsis thaliana* calli grown in presence or absence of Pi. A.** qPCR analysis of AtVPS13M1 mRNA level in Col0 calli grown 6 days in presence (+Pi) or absence (-Pi) of Pi. The mitochondrial malate deshydrogenase (mMDH) was used as a control. **B.C.** PCR of AtVP13M1 fragment used for Oxford Nanopore Technology (ONT) long-read sequencing and results of the FLAIR analysis performed on replicates 1 and 2 in presence or absence of phosphate. **B.** Gels obtained after PCR for amplification of a 7.6kb fragment. The bands in red square were purified after gel excision, and then used for ONT long-read sequencing. **C.** Number of reads and representative % (in brackets) obtained for the 11 isoforms identified by the FLAIR pipeline that represent at least 1% of the reads.

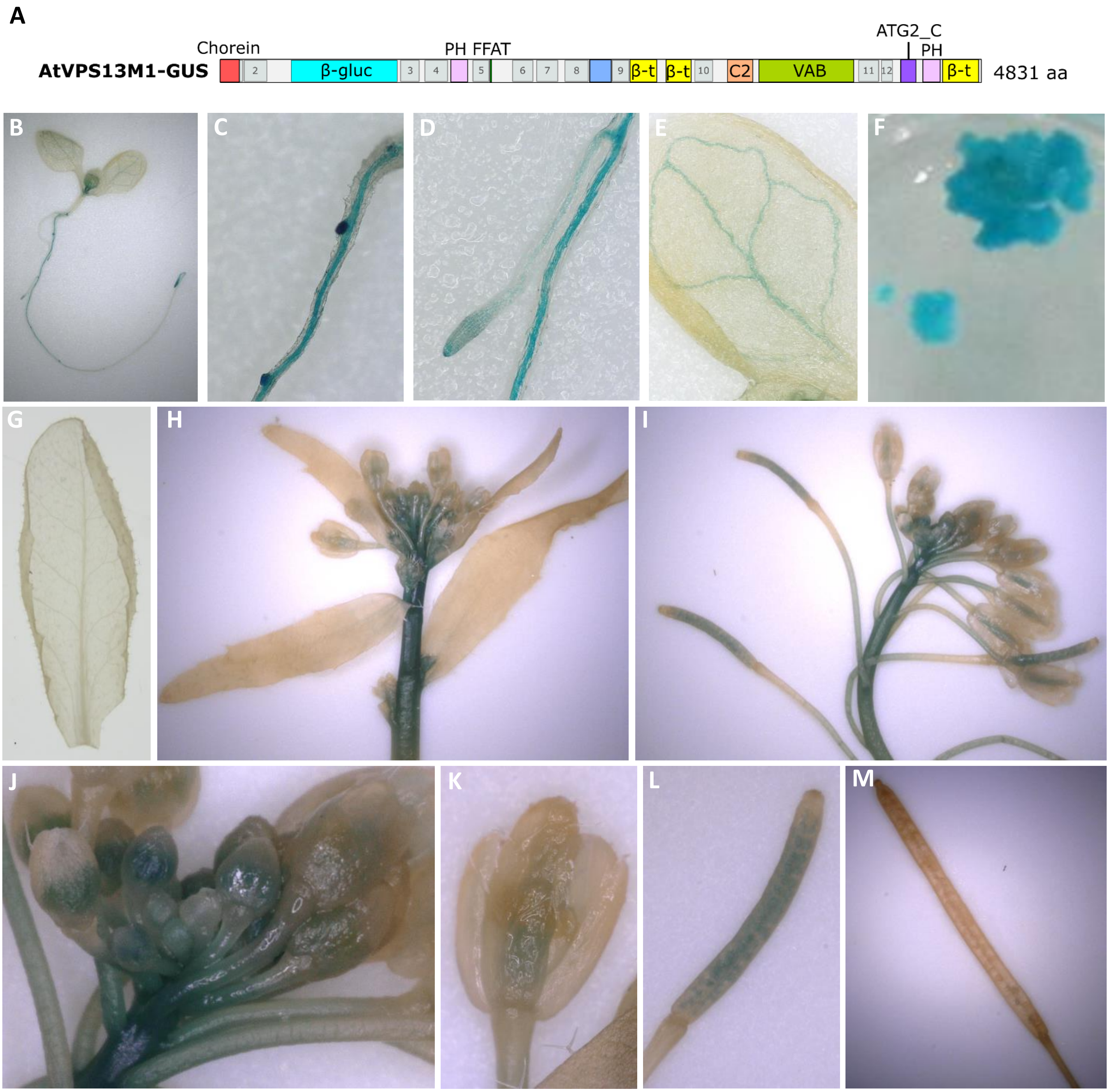


**Figure S2: Analysis of AtVPS13M1 expression in *Arabidopsis thaliana atvps13m1.3* ko line stably expressing AtVPS13M1 fused to β-glucuronidase at position 472 expressed under its native promotor**. **B-F.** Results from stably transformed line n°4 are shown. **G-M.** Results from transformed line n°3 are shown. **A.** Schematic representation of the AtVPS13M1 protein fused to β-glucuronidase (β-gluc) used to analyze AtVPS13M1 expression *in planta*. **B-E.** Analysis of AtVPS13M1 expression by GUS staining in 7-weeks old seedlings grown vertically on plates. **F.** GUS staining of callus. **G-M.** GUS staining of tissues from 7-weeks old plants: leaf (**G**), inflorescences (**H-I**), flower buds (**J**), flowers (**K**), young (**L**) and mature (**M**) siliques.

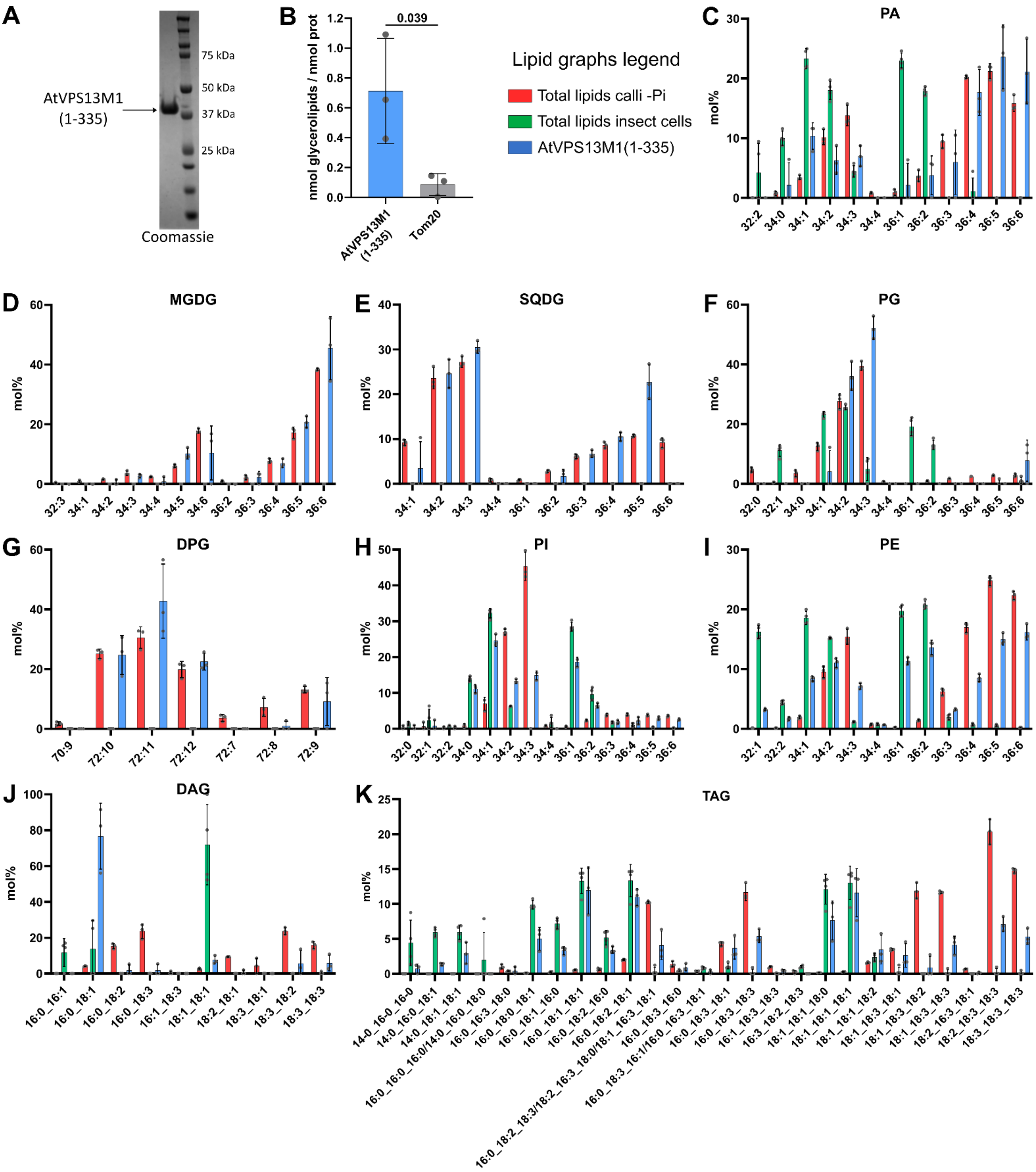


**Figure S3. Analysis of lipids bound to AtVPS13M1(1-335) fragment *in vitro*. A.** Coomassie staining of AtVPS13M1(1-335) fragment expressed and purified from insect cells. **B.** Quantification by mass spectrometry of the total nmol of glycerolipids per nmol of proteins after incubation with calli total lipid extracts. **C-L.** Lipid species found in total lipids from calli grown in –Pi, insect cells and AtVPS13M1(1-335) purified protein incubated with total lipids from calli grown in –Pi. DAG : diacylglycerol, DGDG : digalactosyldiacylglycerol, DPG : diphosphatidyldiacylglycerol, MGDG : monogalactosyldiacylglycerol, PA : phosphatidic acid, PC : phosphatidylcholine, PE : phosphatidylethanolamine, PG : phosphatidylglycerol, PI : phosphatidylinositol, PS : phosphatidylserine, SQDG: sulphoquinovosyldiacylglycerol, TAG: triacylglycerol. Means, standard deviations and individual measures from each biological are represented, N=3.

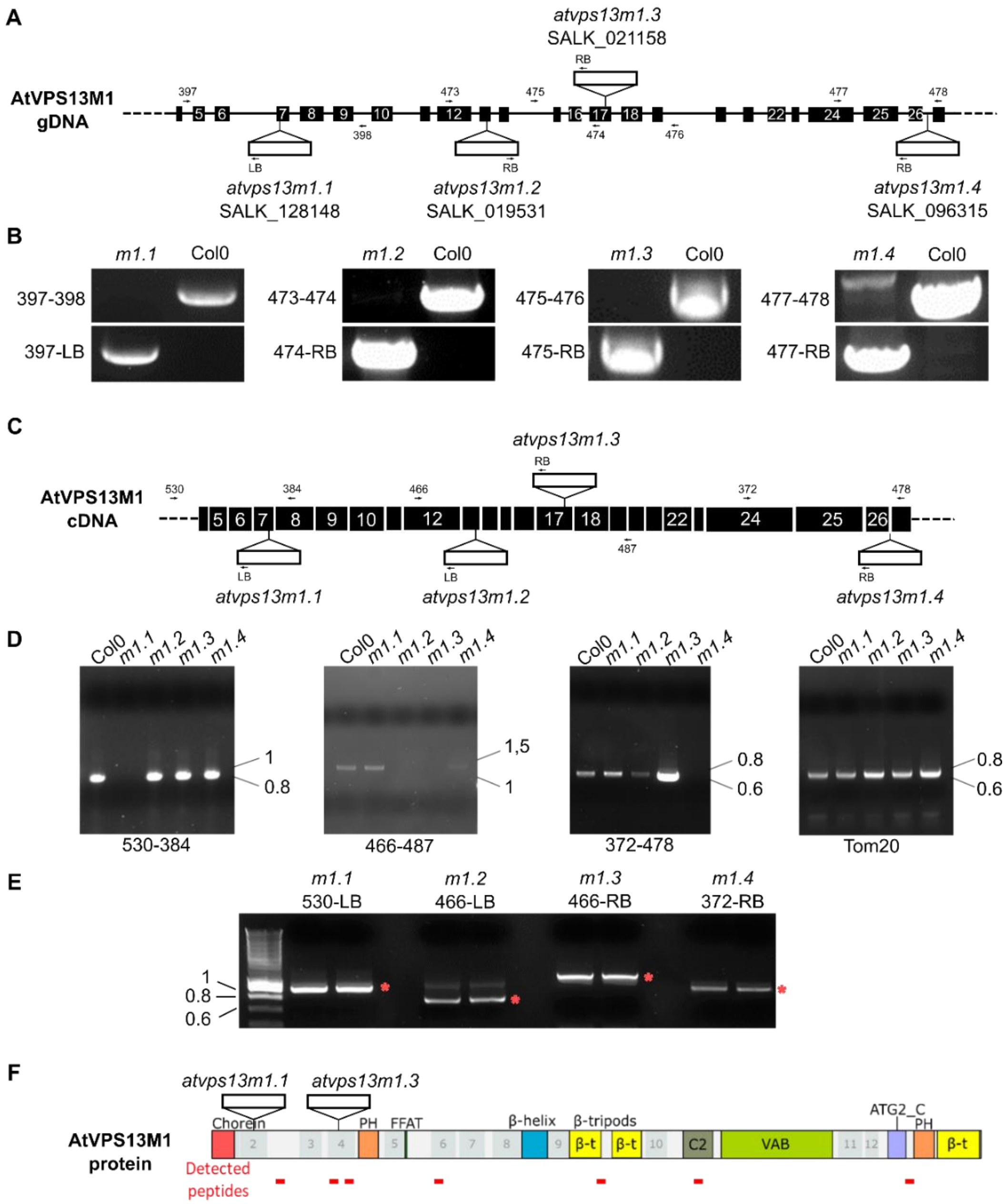


| Line | At4g17140 / AtVPS13M1 | | | At2g31390 / Fructokinase | | | At5g09900 / RPN5A | | |
|---|---|---|---|---|---|---|---|---|---|
| | Repl. 1 | Repl. 2 | Repl. 3 | Repl. 1 | Repl. 2 | Repl. 3 | Repl. 1 | Repl. 2 | Repl. 3 |
| Col0 | 12,9 | 13,2 | 12,5 | 482,6 | 486,8 | 487,3 | 45,1 | 40,9 | 40,4 |
| *atvps13m1.1* | n.d. | n.d. | n.d. | 466,5 | 478,6 | 487,9 | 61,2 | 49,1 | 39,9 |
| *atvps13m1.3* | n.d. | n.d. | n.d. | 476,4 | 479,2 | 484,3 | 51,3 | 48,5 | 43,5 |

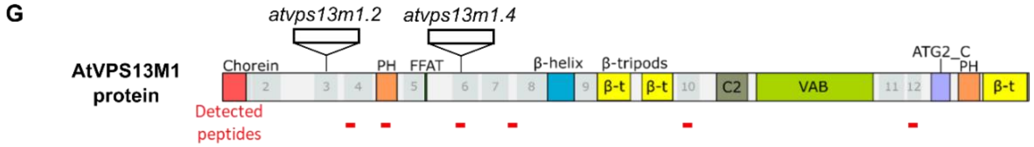


| Line | At4g17140 / AtVPS13M1 | | | AT2G31390 / Fructokinase | | | AT5G09900 / RPN5A | | |
|---|---|---|---|---|---|---|---|---|---|
| | Repl. 1 | Repl. 2 | Repl. 3 | Repl. 1 | Repl. 2 | Repl. 3 | Repl. 1 | Repl. 2 | Repl. 3 |
| Col0 | 7,9 | 8,4 | 13,8 | 485,4 | 491,3 | 482,2 | 42,3 | 36,4 | 45,6 |
| *atvps13m1.2* | nd | nd | nd | 484,6 | 486,1 | 487,5 | 43,1 | 41,6 | 40,2 |
| *atvps13m1.4* | nd | nd | nd | 502,4 | 494,4 | 501,4 | 25,4 | 33,3 | 26,3 |

**Figure S4. Analyses of AtVPS13M1 expression in *atvps13m1* T-DNA insertion lines. A.** Representation of the position of T-DNAs in the four lines analyzed in this study. Only the genomic region from exon 4 to 27 is represented for clarity. **B.** PCR validation of T-DNA insertion and homozygosity for the four *atvps13m1* lines. All lines are homozygous. **C.** Representation of the expected T-DNA insertion positions in AtVPS13M1 mRNA. **D-E.** PCR validation for the presence of the T-DNA in the AtVPS13M1 transcripts. **F-G.** Analysis of AtVPS13M1 protein expression in the different T-DNA insertion lines by MS-based targeted proteomics. Positions of detected peptides along the protein sequence are indicated by red bars. Peptides specific for fructokinase and isoforms of RPN5 were analyzed as controls. For each protein in each sample, abundance (given in $10^6$ arbitrary units) was computed as the sum of specific peptide abundances (Table S1), normalized by the mean of control protein abundances across all samples. *atvps13m1.1* and *m1.3* (**F**) and *m1.2* and *m1.4* (**G**) were analyzed in two different sets of experiments with different peptides (Table S1). Nd: not detected.

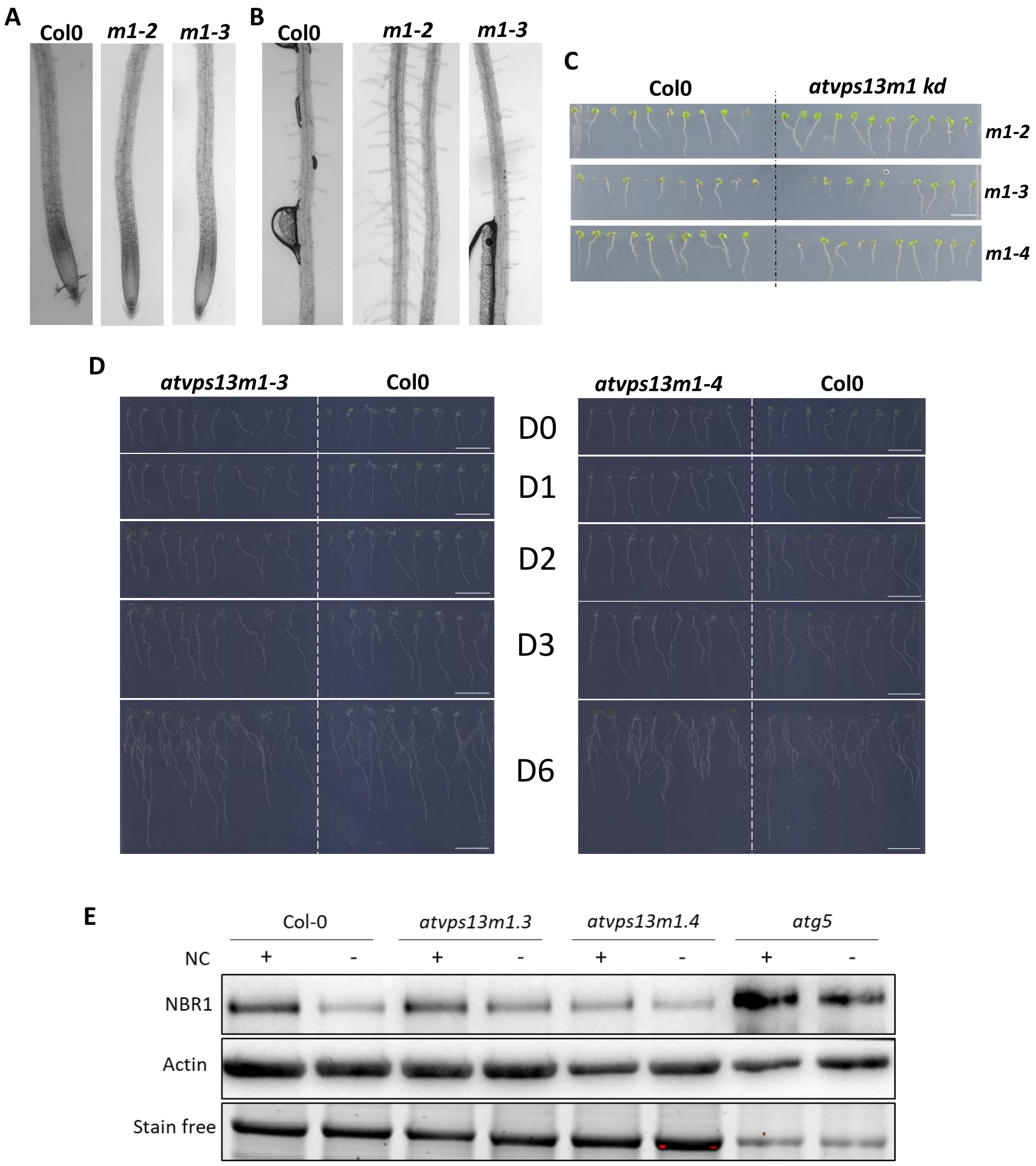


**Figure S5: Phenotype analysis of *atvps13m1* mutant plants. A/B.** Root morphology (**A**) and root hairs analysis (**B**) in 7 days-old seedlings of Col0 and *atvps13m1-2* and *m1-3* mutants. **C.** Representative images of 7-days old seedlings of Col0 and *atvps13m1* kd grown for 7 days in presence of 1 mM phosphate. **D.** Representative series of images of seedlings of Col0, *atvps13m1-3 and m1-4* kd grown for 4 days in 1 mM Pi and transferred in low phosphate (5 µM) (D=0). Growth was followed after 1 (D1), 2 (D2), 3 (D3) and 6 (D6) days after transfer. Scale bar: 1 mm **E.** Representative result of NBR1 protein level analysis by western blot in Col0, *atvps13m1-3*, *atvps13m1-4* and *atg5* mutants. Plants were grown 7 days on plate and then transfered and incubated 6h in liquid media with or without nitrogen en carbon (NC). Actin and total protein staining using Stain free were used as loading controls.

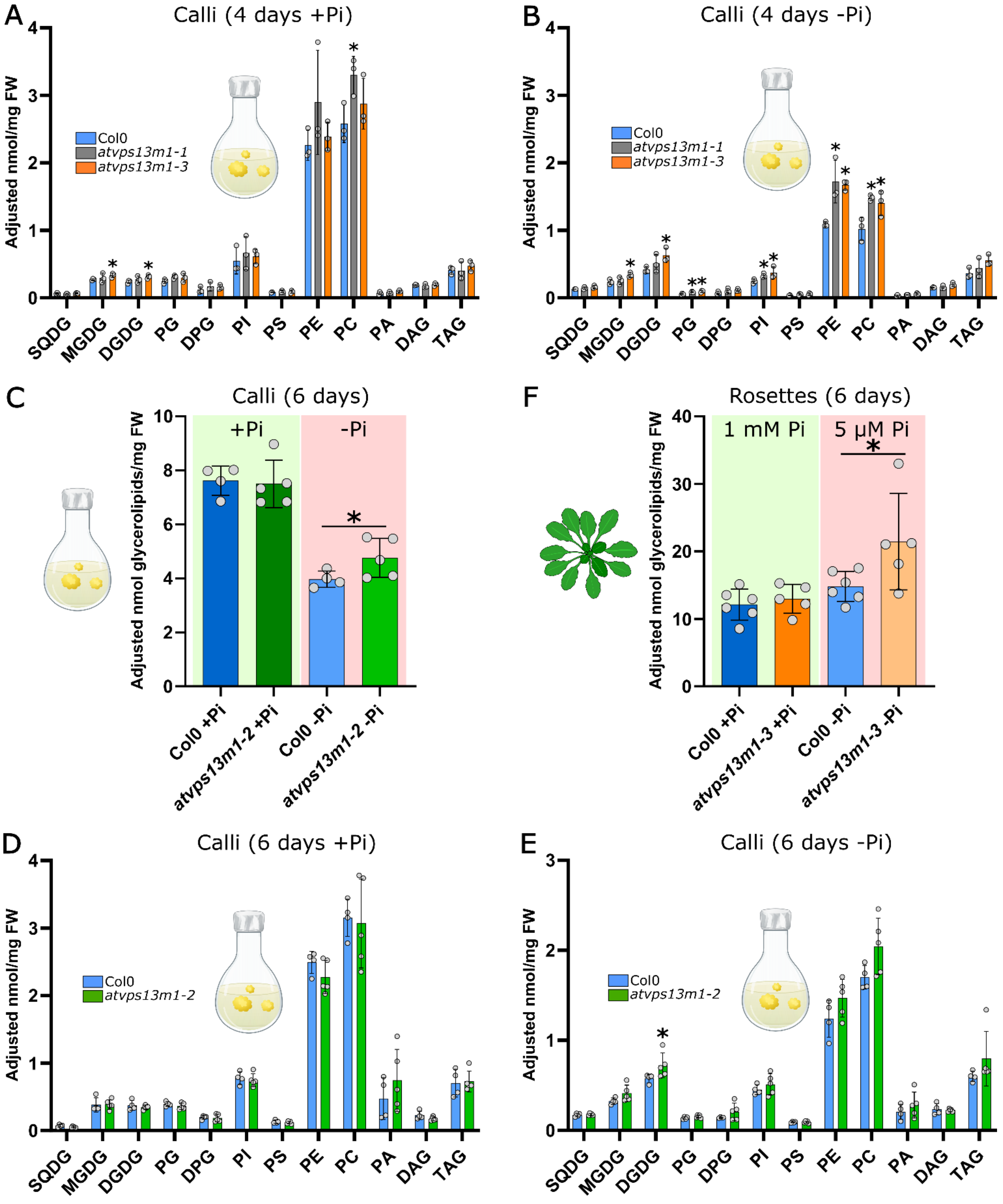


**Figure S6. Lipid analysis in wild type (Col0), *atvps13m1-2* and *m1-3* calli or rosettes grown in presence and absence of phosphate (Pi). A-B**. Lipidomic analyses of Col0 and *atvps13m1-1* and *m1-3* mutants grown 4 days in presence (+Pi) or absence (-Pi) of Pi. Results are expressed in adjusted nmol of each lipid species per mg of calli fresh weight. **C.** Total glycerolipid contents of Col0 and *atvps13m1-2* mutant calli grown for 6 days in presence (+Pi) or absence (-Pi) of Pi. **D-E.** Lipidomic analyses of Col0 and *atvps13m1-2* mutant grown 6 days in presence (**D**) or absence (**E**) of Pi (see material and methods). **F.** Total glycerolipid contents of Col0 and *atvps13m1-3* mutant rosettes grown for 4 days in 1 mM Pi and transferred for 6 days in 1 mM or 5 µM Pi. Statistical analyses were performed using non-parametric Mann-Whitney test. * p-value ≤ 0,05. For all experiments, means, standard deviations and individual measures from each independent biological replicate are represented, N=3-5. DAG : diacylglycerol, DGDG : digalactosyldiacylglycerol, DPG: diphosphatidyldiacylglycerol, MGDG : monogalactosyldiacylglycerol, PA : phosphatidic acid, PC : phosphatidylcholine, PE : phosphatidylethanolamine, PG : phosphatidylglycerol, PI : phosphatidylinositol, PS : phosphatidylserine, SQDG: sulphoquinovosyldiacylglycerol, TAG: triacylglycerol.

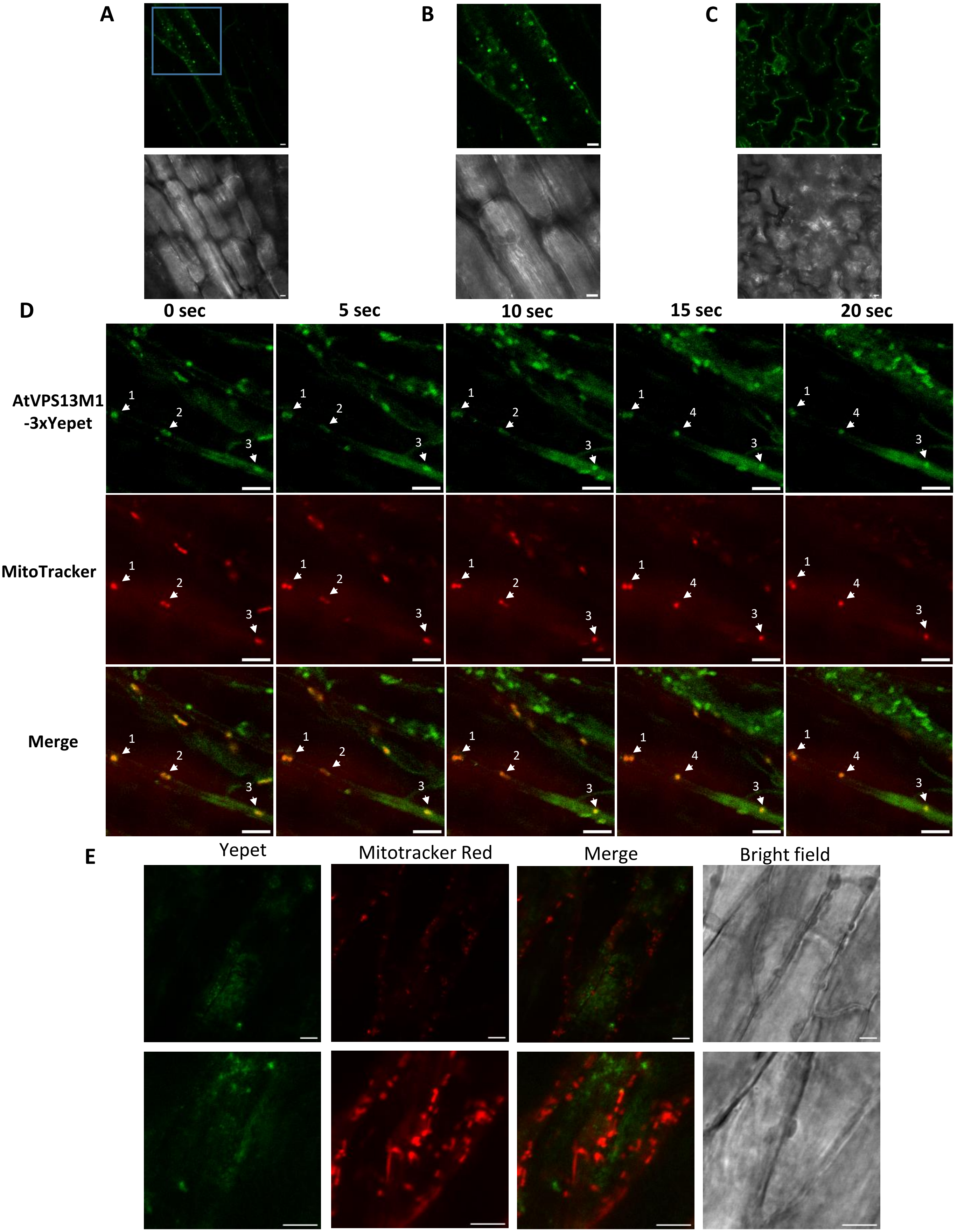


**Figure S7: AtVPS13M1 subcellular localization in 6 weeks-old leaves from *Arabidopsis thaliana atvps13m1-3* mutant stably expressing AtVPS13M1 fused to 3xYepet at position 472 expressed under its native promotor**. **A-C**. Confocal images from stably transformed AtVPS13M1-3xYepet line n°10**. A-B.** Localization in leaves vascular tissue showing a ring-like structure pattern (magnification in **B**). **C**. Localization in epidermal cells. **D.** Time-lapse imaging of AtVPS13M1-3xYepet and mitochondria labeled with Mitotracker$^{TM}$ Red in line n°6. Same mitochondria labeled with AtVPS13M1-3xYepet observed at different time series are labeled by a number. **E.** Yepet fluorescence background obtained in some Col0 leaves vacsular tissues. The background signals do not co-localized with the Mitotracker$^{TM}$ Red. Scale bar: 5µm.

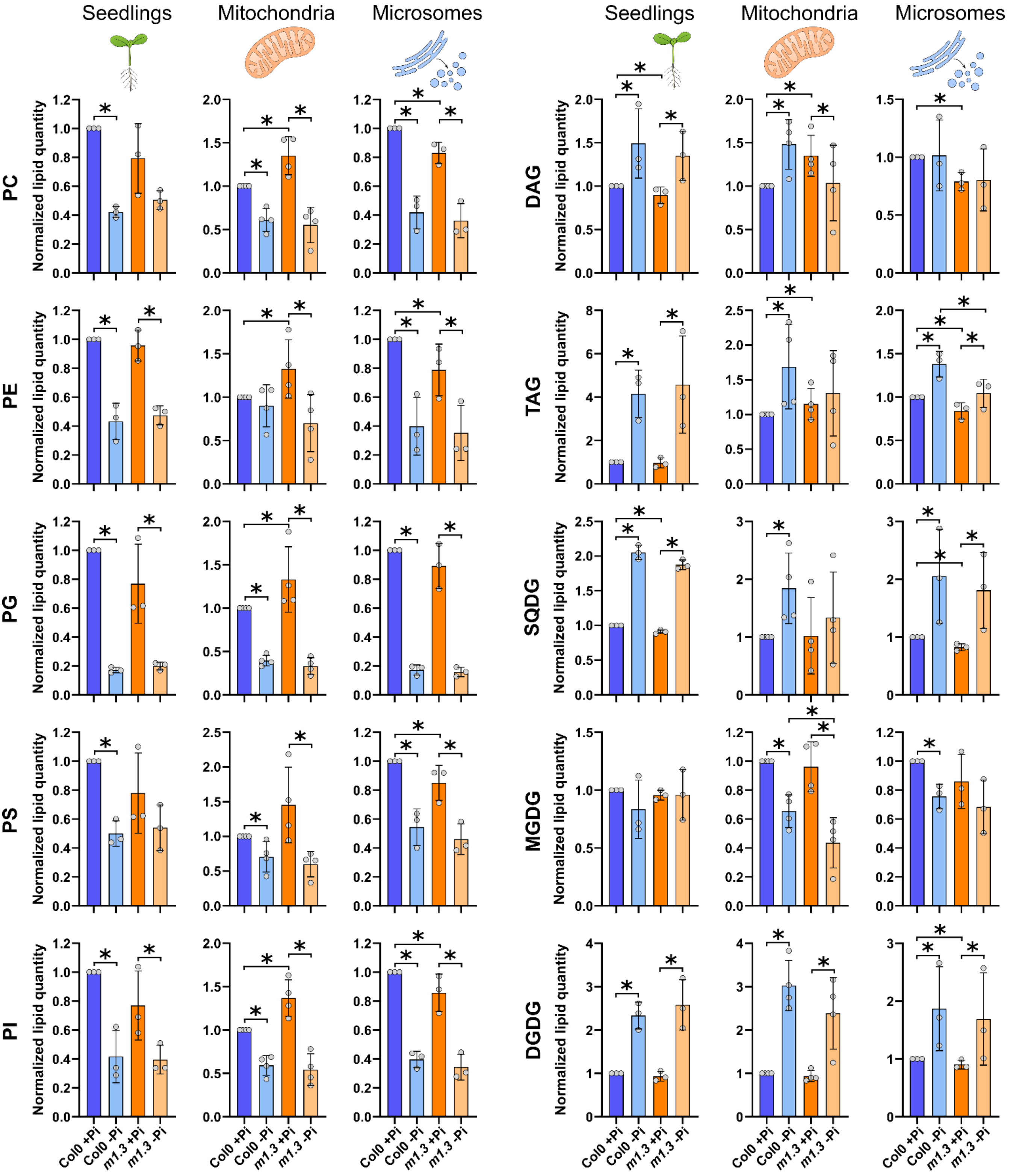


**Figure S8. Lipidomic analysis of wild type (Col0) and *atvps13m1-3* seedlings, mitochondria and microsomes isolated from seedlings grown in presence (+Pi) or absence (-Pi) of phosphate (Pi).** Comparison of the normalized lipid quantity to Col0_+Pi for total glycerolipids or isolated lipid classes from Col0 and *atvps13m1-3* mutant seedlings, mitochondria and microsomes obtained in +Pi or -Pi culture conditions. For seedlings, lipids were quantified in nmol per mg of fresh weight while for mitochondrial and microsomal, lipids were quantified in nmol per mg of protein content (see Material and Methods). On each graph, the means, standard deviations and individual measures from each biological replicate are represented. Statistical analyses were performed using Mann-Whitney tests with a p-value ≤ 0,05 (*).

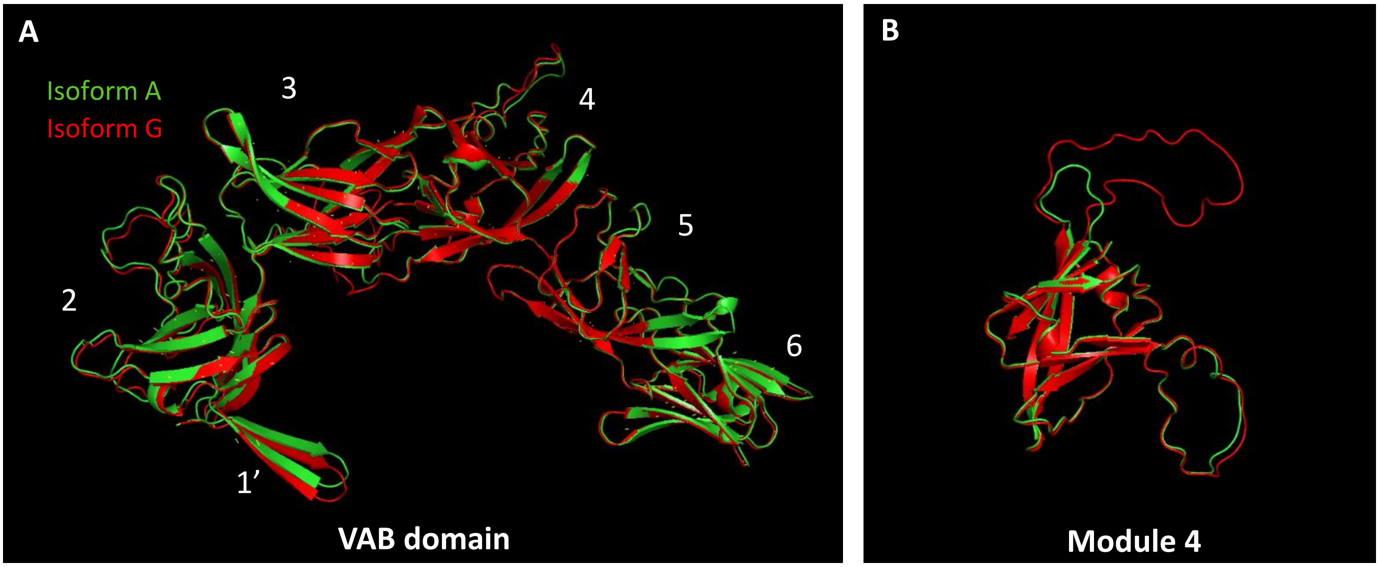


**Figure S9. Alignment of VAB structural prediction of AtVPS13M1 isoform A (in green) and G (in red). A.** The whole VAB domain is presented and the 21 additional amino acids in isoform G did not impact the global folding of the AtVPS13M1 VAB domain. The six repeated modules constituting the VAB domain are annotated. **B.** The 21 additional amino acids in isoform G fold into an longer unfolded loop at the end of the module 4 of the VAB domain. Structure prediction were performed using AlphaFold2 and visualized and aligned using PyMol.
